# Single-cell analysis of innate spinal cord regeneration identifies intersecting modes of neuronal repair

**DOI:** 10.1101/2023.05.19.541505

**Authors:** Vishnu Muraleedharan Saraswathy, Lili Zhou, Mayssa H. Mokalled

## Abstract

Adult zebrafish have an innate ability to recover from severe spinal cord injury. Here, we report a comprehensive single nuclear RNA sequencing atlas that spans 6 weeks of regeneration. We identify cooperative roles for adult neurogenesis and neuronal plasticity during spinal cord repair. Neurogenesis of glutamatergic and GABAergic neurons restores the excitatory/inhibitory balance after injury. In addition, a transient population of injury-responsive neurons (iNeurons) show elevated plasticity between 1 and 3 weeks post-injury. We found iNeurons are injury-surviving neurons that acquire a neuroblast-like gene expression signature after injury. CRISPR/Cas9 mutagenesis showed iNeurons are required for functional recovery and employ vesicular trafficking as an essential mechanism that underlies neuronal plasticity. This study provides a comprehensive resource of the cells and mechanisms that direct spinal cord regeneration and establishes zebrafish as a model of plasticity-driven neural repair.

## INTRODUCTION

Mammalian spinal cord injuries (SCI) elicit complex multi-cellular responses that impede regeneration and cause permanent functional deficits in mammals (Brennan and Popovich, 2018; Fouad et al., 2021; He and Jin, 2016; Milich et al., 2019; Sofroniew, 2018; Tran et al., 2022). Anti-regenerative neuron-extrinsic factors comprise chronic inflammation, fibrotic scarring, demyelination and the acquisition of a regeneration restricting extracellular milieu. These injury complications exacerbate the inherently limited ability of the mammalian spinal cord (SC) to replenish lost neurons via adult neurogenesis or to regrow lesioned axon tracts. Consequently, even the most groundbreaking regenerative therapies targeting select cell types or individual molecules have only yielded modest improvement in cellular and functional outcomes (Park et al., 2008; Sofroniew, 2018). SCI studies have since pursued combinatorial strategies as a more promising therapeutic avenue (Anderson et al., 2018; DePaul et al., 2017; Nakamura et al., 2021; Squair et al., 2023). We propose that comprehensive and simultaneous examination of neuronal and non-neuronal cells after SCI is fundamental to understanding and manipulating the multi-cellular complexities of neural injuries.

Unlike mammals, adult zebrafish have an innate ability to spontaneously recover from severe SCI. Following complete transection of SC tissues, zebrafish reverse paralysis and regain swim function within 6 to 8 weeks of injury (Burris et al., 2021; Mokalled et al., 2016). Pro-regenerative injury responses involving immune and progenitor cells, neurons and glia, cooperate to achieve spontaneous and efficient repair in zebrafish (Briona et al., 2015; Cavone et al., 2021; Klatt Shaw et al., 2021; Kuscha et al., 2012; Reimer et al., 2008; Saraswathy et al., 2022; Vandestadt et al., 2021). Early after SCI, potent populations of progenitor cells, including central canal-surrounding ependymo-radial glial cells (ERGs), are activated to replenish lost neurons and glia (Briona *et al*., 2015; Klatt Shaw *et al*., 2021; Reimer *et al*., 2008; Saraswathy *et al*., 2022). Newly differentiated motor neurons and interneurons populate the regenerate tissue, as pre-existing neurons regrow axons across lesioned tissues (Kuscha *et al*., 2012; Reimer *et al*., 2008; Saraswathy *et al*., 2022). Though less studied, glial cells are thought to enact instrumental pro-regenerative responses throughout the course of regeneration (Goldshmit et al., 2012; Klatt Shaw *et al*., 2021; Mokalled *et al*., 2016). However, a holistic understanding of the cellular interactions that coordinate the pro-regenerative responses to direct SC regeneration in zebrafish is to be acquired.

The advent of single-cell transcriptomics provided the tools to achieve a refined understanding of molecular SCI responses across species and cell types. Multiple single-cell RNA sequencing (scRNA-seq) atlases have been generated for mouse SCI, rat SCI or human SC tissues (Matson et al., 2022; Milich et al., 2021; Rodrigo Albors et al., 2023; Yadav et al., 2023; Yao et al., 2022). Single-cell studies from mice revealed new insights into macroglial cell-cell interactions after injury, revealed a rare population of lumbar spinocerebellar neurons that elicit a regeneration signature, or characterized the responses of astrocytes after injury (Hou et al., 2022; Matson *et al*., 2022; Milich *et al*., 2021; Rodrigo Albors *et al*., 2023). However, single-cell studies from zebrafish SC tissues have been limited to isolated immune or progenitor motor neuron cells and to larval stages (Cavone *et al*., 2021; Scott et al., 2021). Thus, a complete resource of the regenerative cells and mechanisms from adult zebrafish is required to develop a molecular understanding of the cell identities that enable or limit spontaneous plasticity and regeneration.

The fundamental principles that determine or limit regenerative capacity across species have eluded scientists for ages. While pro-regenerative injury responses are overwhelmed by anti-regenerative complications in mammals, zebrafish cells exhibit increased potency and exclusively pro-regenerative signatures after SCI. Molecular and cellular studies of select cell types or regenerative pathways revealed key insights into the cellular contributions and molecular signatures associated with elevated regenerative capacity. Specifically, while ependymal cells elicit limited stem cell potential in adult mice (Ren et al., 2017; Shah et al., 2018), zebrafish ERGs retain radial glial features and contribute to neurogenesis and gliogenesis after SCI (Briona *et al*., 2015; Klatt Shaw *et al*., 2021; Reimer *et al*., 2008; Saraswathy *et al*., 2022). The disparities in neuron progenitor capacities between zebrafish and mammals yielded an assumption that neurogenesis-based neural repair is unachievable in mammals and directed the community’s efforts toward plasticity-based neural repair strategies. However, while zebrafish has been an established model of neuron regeneration, it remains unclear whether zebrafish could contribute insights and applications into plasticity-driven repair mechanisms. Thus, how and why neuronal injury responses differ between zebrafish and mammals require comprehensive molecular investigation and cross-species comparisons.

This study presents an atlas of the dynamic responses across major spinal cell types during early, intermediate and late stages of regeneration in adult zebrafish. Single nuclear RNA sequencing (snRNA-seq) was performed at 0, 1, 3 and 6 weeks post-injury. Neurons elicit elevated signaling activity relative to the dozens of cell types that respond to injury. While SCI disrupts the excitatory/inhibitory neuron balance, sequential neurogenesis of excitatory and inhibitory neurons restores the homeostatic neuronal landscape at late stages of regeneration. In addition to regenerating new neurons, a transient regenerative signature emerges in a population of injury-responsive neurons (iNeurons) between 1 and 3 weeks post-injury. EdU labeling showed iNeurons are injury surviving neurons that acquire a neuroblast-like transcriptional signature after injury. iNeuron markers genes are required for functional SC repair, and dynamic vesicular trafficking is a central mechanism that promotes spontaneous neuronal plasticity. This study identifies multi-layered modes of regenerative neurogenesis and neuronal plasticity during innate SC regeneration, and establishes zebrafish as a platform to identify and manipulate regeneration- and plasticity-based modes of neural repair.

## RESULTS

### Molecular identification and temporal dynamics of spinal cell types after injury

To determine the transcriptional identities of the cells that directs successful SC regeneration, we performed complete SC transections on adult zebrafish and dissected 3 mm sections of SC tissues surrounding the lesion site at 1, 3, and 6 weeks post-injury (wpi) for nuclear isolation. Corresponding tissue sections were collected from uninjured controls. SC tissues were pooled from 50 animals per time point, and 2 pools of independent biological replicates were analyzed for each time point. Our dataset spans key regenerative windows including early injury-induced signals at 1 wpi, neuronal and glial regeneration at 3 wpi, and cellular remodeling at late stages of regeneration at 6 wpi (Mokalled *et al*., 2016). Isolated nuclei were sequenced using 10× genomics platform (3’ v3.1 chemistry) and aligned to zebrafish genome GRCz11 with improved zebrafish transcriptome annotation (Lawson et al., 2020; Matson et al., 2018). Nuclei were subsequently filtered using the Decontx and DoubletFinder packages to exclude droplets that include doublet nuclei or a high proportion of ambient mRNA, respectively (McGinnis et al., 2019; Yang et al., 2020). A second round of filtering is performed by thresholding the covariates such as number of counts, genes and fraction of counts from mitochondrial genes per barcode. A total of 58,973 nuclei was obtained for downstream analysis using the Seurat package (Fig. 1A and S1A-C).

**Figure 1.**
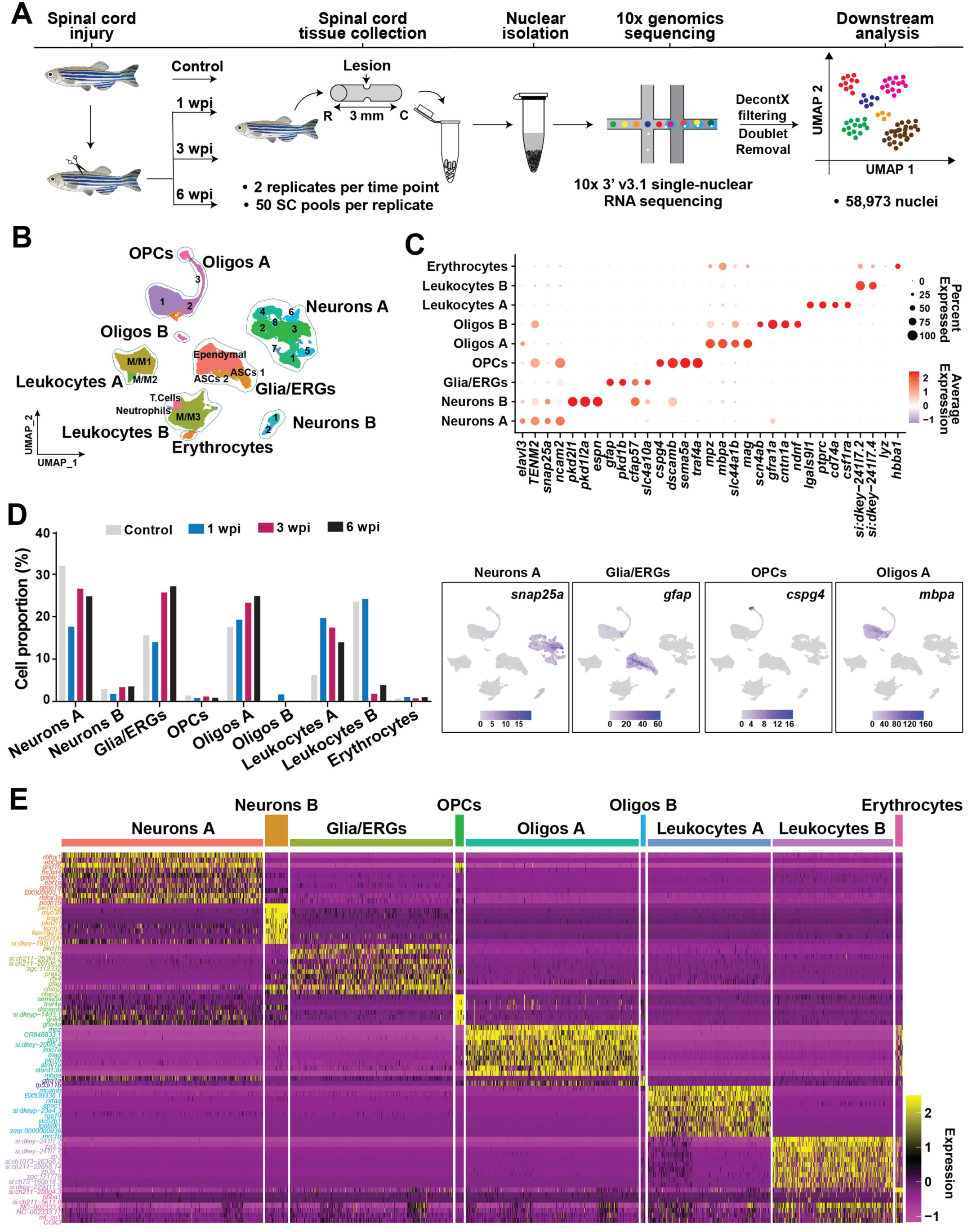
Transcriptomic profiling of innate SC repair in adult zebrafish. **(A)** Experimental pipeline to generate a single-cell atlas of zebrafish cells after SCI. SC tissue collection, nuclear isolation and single nuclear RNA-seq (snRNA-seq) were performed on wild-type fish at 1, 3 and 6 wpi. Uninjured SC nuclei were used as 0 wpi controls. 10x genomics sequencing with v3.1 chemistry was performed. Two biological replicates were used at each time point, and SC tissues from 50 animals were pooled into a single biological replicate. **(B)** Merged UMAP representation of the complete dataset. Two biological replicates, 4 time points and 58,973 cells were clustered into major spinal cell populations including Neurons A and B, glia/ependymo-radial glial cells (glia/ERGs), oligodendrocyte precursor cells (OPCs), oligodendrocytes A and B, Leukocytes A and B, and Erythrocytes. **(C)** Marker gene expression in the major cell types identified after SCI. Dot plot shows canonical marker genes are enriched in their respective cell types identified. Dot colors and diameters represent average gene expression and percent cells with at least one UMI detected per gene, respectively. Feature plots depicting the distributions of canonical marker genes corresponding to the major cell populations are shown. **(D)** Distribution of major cell types during SC regeneration. For each time point, cell proportions were normalized to the total number of cells present at that time point. **(E)** Heatmap showing top 10 differentially expressed (DE) genes in the major cell populations identified by snRNA-seq. Yellow and magenta represent highly enriched and low-expression genes, respectively.

Clustering of SC nuclei revealed 24 cell clusters with distinct molecular identities and temporal dynamics (Fig. S1D and Table S3). Postulating that previously established cell type classifiers are likely biased toward cell identities present in the mammalian nervous system or in developing zebrafish, we optimized cell type identification by generating a custom-assembled <u>V</u>ertebrate C<u>N</u>S <u>M</u>arker (VNM) database. VNM compiled cell type markers commonly present in the central nervous system from over 12 datasets and from vertebrate species including zebrafish, mice and humans (Table S1) (Baek et al., 2019; Cavone *et al*., 2021; Guillemot, 2007; Haring et al., 2018; Hayashi et al., 2018; Hernandez-Miranda et al., 2017; Lu et al., 2015; Milich *et al*., 2021; Rosenberg et al., 2018; Rougeot et al., 2019; Sathyamurthy et al., 2018; Tambalo et al., 2020; Tang et al., 2017; Zeisel et al., 2018; Zhang et al., 2014). We then cross-referenced differentially expressed (DE) markers for each cluster with our VNM database (Table S2 and Fig. S1E). Scoring matrix heatmap representation of this analysis identified 6 major cell types comprising neurons, glia/ERGs, oligodendrocyte precursor cells (OPCs), oligodendrocytes, leukocytes and erythrocytes. Two distinct cell populations were identified for neurons (Neurons A and B), oligodendrocytes (Oligos A and B) and Leukocytes (Leukocytes A and B). Additional subclusters within each major cell population were further identified using the prediction from scoring matrix heatmap (Fig. 1B and S1E). VNM-based cell type classification was further confirmed by evaluating the expression of canonical marker genes for each cluster (Fig. 1C). *elavl3* and *snap25a* were enriched in Neurons A and B, *gfap* and *slc4a10a* in glia/ERGs, *cspg4* and *sema5a* in OPCs, *mpz* and *mbpa* in oligodendrocytes A, *csfr1a* and *ptprc* in leukocytes A, *hbba1* in erythrocytes (Fig. 1C and S1G). Although canonical cell markers were enriched in their respective clusters, the proportions of cells expressing these markers in leukocytes A, glia/ERGs and erythrocytes were relatively small. On the other hand, gene expression analysis of Neurons B, Oligodendrocytes B and Leukocytes B did not show enrichment for classical markers. We conclude that our VNM database is likely still biased toward commonly used mammalian cell markers that are not necessarily the best markers for adult zebrafish, and that our single-cell atlas provides a platform to identify improved CNS markers for adult zebrafish cells. To test this hypothesis, we examined whether our snRNA-seq dataset identified improved cell-specific DE markers in adult zebrafish SCs (Fig. 1E). We found that Glia/ERG cells expressed the cilia-associated *cfap57* gene, indicative of the ciliated nature of the SC central canal-lining ERGs. Neurons B expressed *pkd2l1* and *pkd1l2a*, which are known markers of cerebrospinal fluid-contacting neurons (Sternberg et al., 2018). Oligodendrocytes B expressed *scn4ab* and *gfra1a*. Leukocytes A were enriched for *mrc1b* and *slco2b1* along with the cell adhesion molecule coding *mcamb* gene, whereas Leukocytes B expressed unknown genes such as *si:dkey-241l7.2* and *si:dkey-241l7.4*. These studies indicated that extrapolating mammalian CNS or developmental zebrafish markers in adult zebrafish requires validation, and provided a platform to identify new and improved CNS cell markers in adult zebrafish.

To examine the temporal dynamics of SC cells during regeneration, we determined the numbers and relative proportions of nuclei within each coarse cluster in the integrated dataset (Fig. 1D and S1F). Neurons A, which comprised 32% of the nuclei harvested from uninjured SCs, comprised 18% of total nuclei at 1 wpi and recovered to 25% at 6 wpi. Neurons B represented a smaller proportion of neurons and showed little variation during the course of regeneration. The proportions of leukocytes A increased from 6% of control nuclei prior to injury to 20% at 1 wpi and gradually decreased to 14% at 6 wpi. We observed an expansion in the relative abundance of glia/ERGs between 1 and 3 wpi, whereas OPCs maintained similar proportions at all time points. Oligodendrocytes A, which accounted for 18% of uninjured nuclei, gradually increased to 25% of total nuclei at 6 wpi. In contrast, Oligodendrocytes B cells were transiently present at 1 wpi and comprised 2% of total nuclei at that time point. These studies provided a comprehensive resource of the cellular architecture that directs successful SC regeneration and revealed a gradual recovery in neuronal proportions during innate SC repair.

### Neurons are active signaling hubs during spinal cord regeneration

To infer molecular modes of intercellular communication during SC regeneration, we surveyed signaling pathways that are differentially active between 0 and 6 wpi (Fig. 2 and S2). We first converted our dataset of zebrafish genes to their human orthologues using the BioMart package from Ensembl. The R package “CellChat” was then used to quantitatively analyze global intercellular communication networks based on the expression of ligands, receptors and cofactors (Jin et al., 2021). CellChat computes the communication probabilities of ligand-receptor interactions associated with each signaling pathway, and integrates the cumulative probabilities of all signaling pathways that are outgoing or received (incoming) by each cell population. In control SC tissues, the cumulative strengths of all signaling pathways outgoing from Neurons A and B were the highest relative to other cell types, indicating active neuronal signaling under homeostatic conditions (Fig. 2A-B). After SCI, outgoing signaling increased in most cell types, indicating tissue-wide responses are activated after injury (Fig. 2A-B). Notably, the strengths of signaling pathways outgoing from Neurons A and B continued to be elevated across all time points and increased more than 2.5 fold between 0 and 6 wpi (Fig. 2A-B). On the other hand, the cumulative strengths of all incoming signaling pathways received by OPCs were the highest among spinal cell types after SCI (Fig. S2A and 2B). Incoming signaling received by Neurons A doubled between 0 and 6 wpi (Fig. 2B). Mapping of ligand-receptor interactions predicted cell-cell signaling interactions are highest between neurons and OPCs at 6 wpi (Fig. 2C). Confirming CellChat analysis, trajectory inference showed OPCs and neurons have elevated transcriptional activity during SC regeneration (Gulati et al., 2020) (Fig. S2B). These results indicated neurons and OPCs have elevated and dynamic transcriptional and signaling activities during zebrafish SC regeneration.

**Figure 2.**
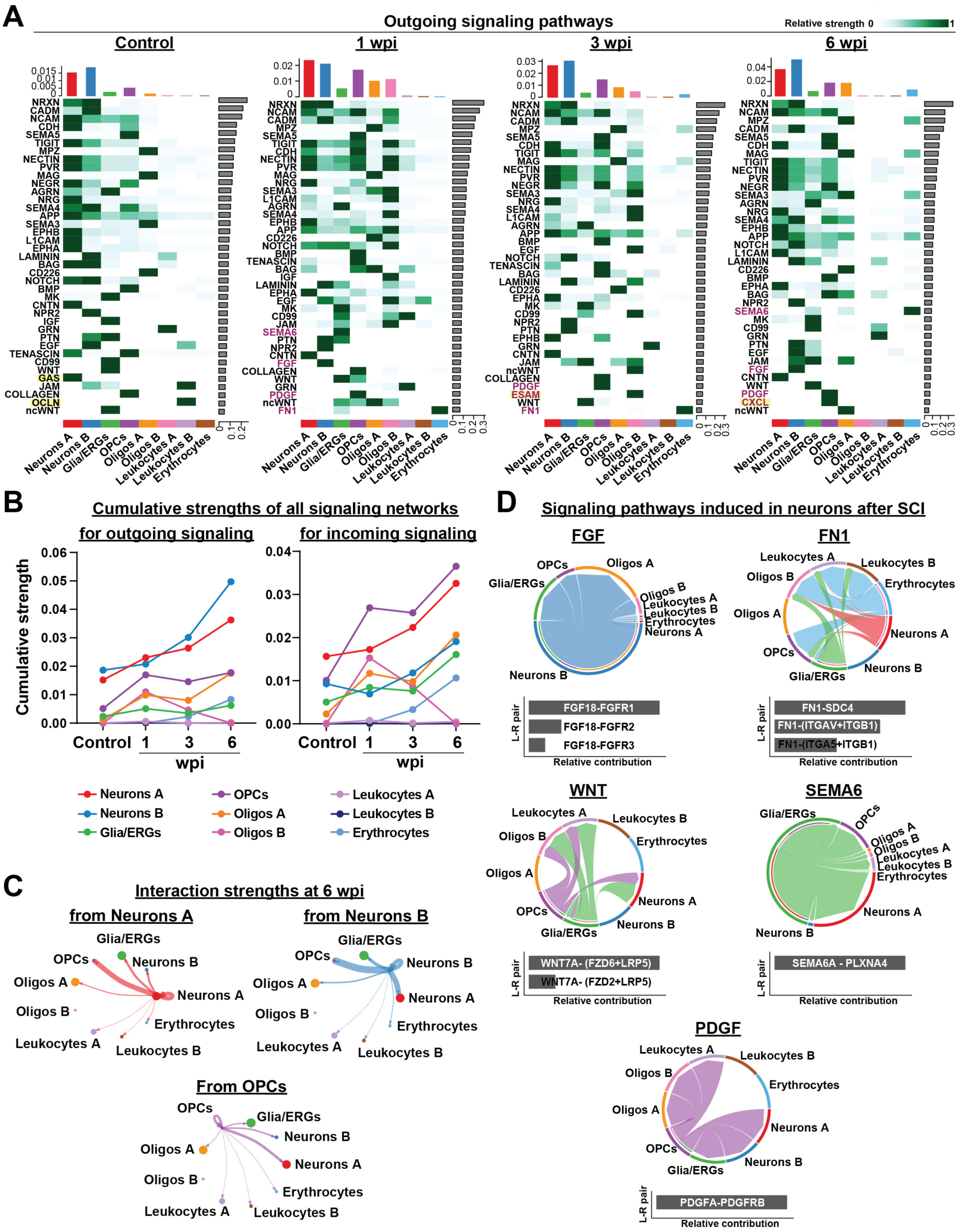
Cell-cell interaction networks during SC regeneration. **(A)** Relative strengths of outgoing signaling pathways in major spinal cell populations at 0, 1, 3 and 6 wpi. Bar graphs at the top of each heatmap show cumulative signaling strengths per cell population. Bar graphs at the right of each heatmap show cumulative signaling strength per pathway. Pathways highlighted in yellow are specifically enriched at one time point. Pathways highlighted in magenta are enriched after injury. **(B)** Cumulative strengths of signaling pathways outgoing from or received by major cell types. **(C)** Circle plots represent cell-cell interaction strengths of signaling pathways outgoing from Neurons A, B and OPCs at 6 wpi. Arrow thickness is directly proportional to the cumulative strengths of all predicted interactions between clusters. **(D)** Chord diagrams showing intercellular communication of signaling pathways that are enriched in neurons after SCI. Arc diameters are proportional to cumulative signaling strengths. Bar graphs at the bottom of each chord diagram represent the relative contribution of different ligand-receptor pairs predicted for the corresponding signaling pathway.

### Global and neuron-specific analyses of injury-induced signaling pathways

Global CellChat analysis revealed active signaling pathways in control, 1, 3 and 6 wpi SC tissues. Tracking the strengths of individual signaling pathways across time points indicated dynamic intercellular communication during SC regeneration (Fig. 2A and S2A). In addition to dynamic regulation of signaling pathways present in control SC tissues, 7 pathways were specifically activated at 1, 3 or 6 wpi relative to uninjured controls. Globally-induced signaling pathways included chemokine signaling (CXCL), cell adhesion signaling networks (FN1 and ESAM), extracellular matrix (ECM) and growth factor mediated signaling networks (SEMA6 and PDGF) and developmental signaling pathways (FGF) (Fig. 2A and S2A).

To identify molecular pathways that account for elevated neuron signaling, we analyzed signaling pathways predicted to be outgoing from or received by Neurons A and B (Fig. 2D and S2C). Including both incoming and outgoing signaling, 32 to 35 pathways were predicted to be activated by neurons between 0 and 6 wpi (Fig. S2C). In addition to homeostatic signaling pathways increasing in strengths after injury, 5 injury-induced pathways were either activated or received by neurons after injury (FGF, WNT, PDGF, FN1 and SEMA6) (Fig. 2D). SCI triggered reactivation of growth factor and developmental signaling, substantiating FGF, WNT and PDGF as regulators of neuron survival, axon regrowth and synapse formation (Cardozo et al., 2017; Diez Del Corral and Morales, 2017; Garcia et al., 2018; Goldshmit *et al*., 2012; Guo et al., 2019; He et al., 2018; Shimada et al., 2016; Tome et al., 2023; Tsata et al., 2021; Ye et al., 2021). Specifically, Neurons B are a major source of FGF18, which is predicted to interact with multiple FGFR1/2/3-expressing spinal cells populations. On the other hand, Neurons A are predicted recipients of WNT signaling, whereas both Neurons A and B are at the receiving end of PDGF signaling. Ligand-receptor interactions involve WNT7A-FZD2/6/LRP5 and PDGFA-PDGFRB, whereby Glia/ERG- and OPC-derived WNT7A and PDGFA are received by FZD2/6/LRP5- and PDGFRB-expressing neurons. Highlighting the requirement for extracellular matrix (ECM) remodeling and signaling to enact a pro-regenerative response (Bradbury et al., 2002; Monaghan et al., 2007; Zavvarian et al., 2021), multiple cell types including Neurons A express Fibronectin 1 (FN1), which is predicted to direct strong interactions with SDC4-, ITGAV-, ITGA5- and ITGB1-expressing cells. Conversely, Glia/ERGs are a major source of outgoing SEMA6A signaling, received by PLXNA4-expressing neurons to regulate axon guidance (Carulli et al., 2021; Runker et al., 2008; Squair *et al*., 2023; Syed et al., 2016; Yoshida, 2012). These studies revealed complex multicellular communication networks that coordinate signaling between neurons, glia, and non-neural injury-responding cells during SC repair. Further validation and mechanistic insights into these signaling pathways are required to understand their exact function during zebrafish SC regeneration.

### Neuronal E/I balance is restored during successful spinal cord repair

To explore the cellular dynamics that support neuronal repair and functional recovery after SCI, we examined neuronal cell types and states prior to and after injury. Subcluster analysis identified 30 distinct neuron clusters between 0 and 6 wpi (Fig. 3A, S3A-B and Table S4). Balanced synaptic excitatory and inhibitory (E/I) transmission is a central feature of CNS function as altered E/I balance results in severe behavioral defects in humans (Isaacson and Scanziani, 2011). Despite eliciting features of cellular regeneration, SCI in mouse neonates causes impaired locomotor recovery due to imbalanced E/I transmission (Bertels et al., 2022; Dudek and Sutula, 2007; Gao and Penzes, 2015). With zebrafish recovering swim function after SCI, we postulated that balanced E/I transmission is a hallmark of functional SC repair in zebrafish. To test this hypothesis, we first performed a bioinformatic characterization of the excitatory and inhibitory neurotransmitter properties of the neuron clusters identified in our dataset (Fig. 3B). Gene expression for classical markers of glutamatergic (slc*17a6a, slc17a6b*), GABA/glycinergic (*slc6a5, gad1a, gad1b, gad2*), cholinergic (*chata, slc5a7a*), serotonergic (*slc6a4a, tph2*) and catecholaminergic (*slc6a2*) neurons were analyzed (Horzmann and Freeman, 2016) (Fig. S3C). Neuron subclusters were then grouped as excitatory, inhibitory or ‘Other’ based on their neurotransmitter gene expression profiles (Fig. 3B and S3C-E). The proportions of excitatory (30%) was lower than inhibitory (56%) neurons in uninjured SC tissues (Fig. 3C), resulting in a baseline E/I ratio of 0.5 (Fig. 3D). The profile of excitatory neurons increased to 34% of total neurons at 1 wpi and returned to baseline levels at 6 wpi. Inhibitory neurons did not change in proportions at 1 wpi but gradually increased to 60% at 6 wpi, suggesting a compensatory mechanism for early surge in excitatory neurons in the injured spinal cord (Fig. 3C). *in silico* calculation of the E/I ratio confirmed an imbalance toward an excitatory phenotype at 1 wpi and recovery to baseline by 6 wpi (Fig. 3D).

**Figure 3.**
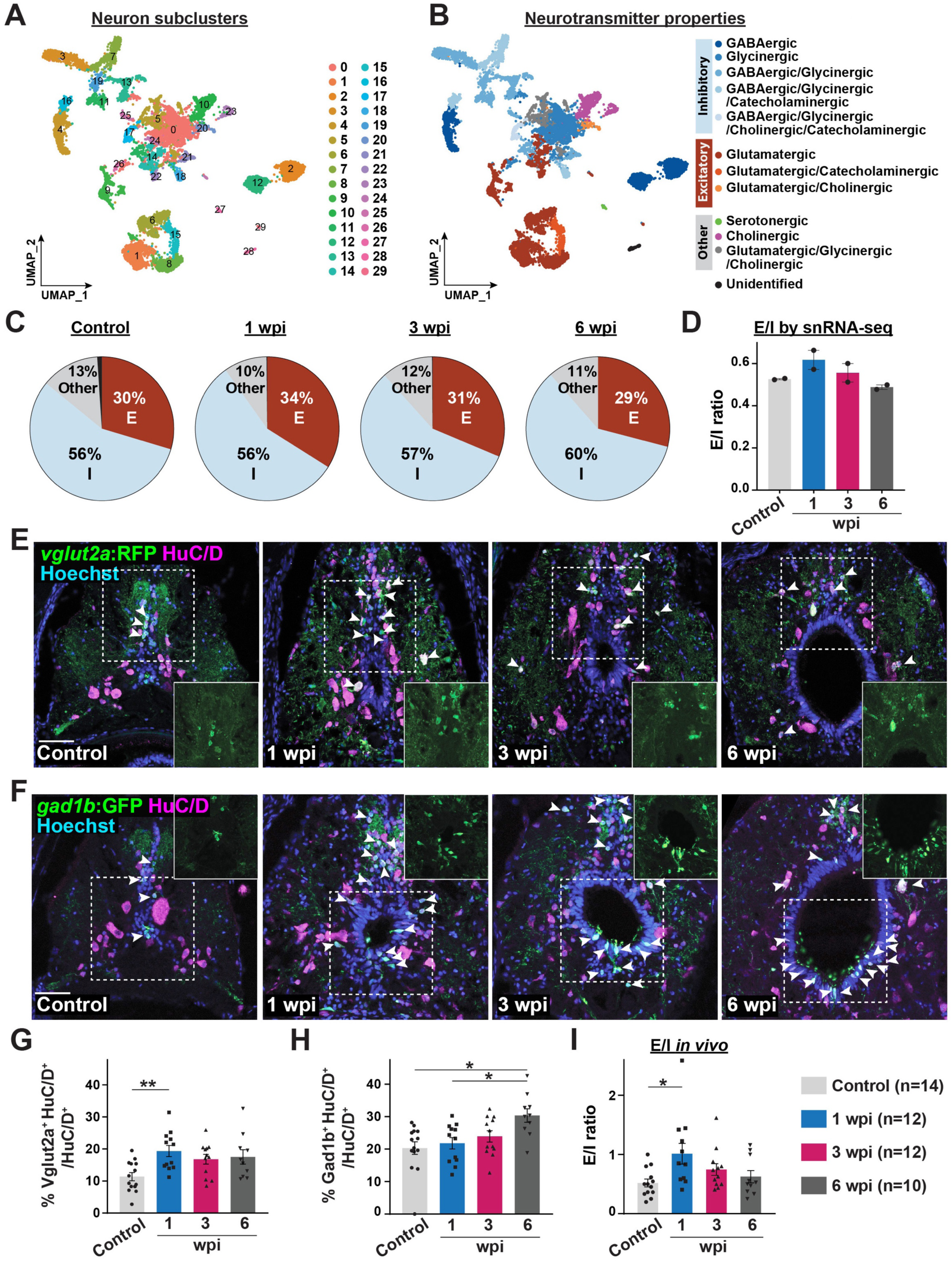
Recovery of excitatory/inhibitory balance during SC regeneration. **(A)** UMAP plot showing 30 different neuron subclusters identified with 0.6 resolution parameter. **(B)** Classification of neuron clusters based on neurotransmitter properties. Canonical neurotransmitter marker genes are used: *slc17a6a* and *slc17a6b* for glutamatergic neurons; *slc6a5, gad1a, gad1b* and *gad2* for GABA/glycinergic neurons; *chata and slc5a7a* for cholinergic neurons; *slc6a4a* and *tph2* for serotonergic neurons and *slc6a2* for catecholaminergic neurons. Clusters with enrichment of at least one of the glutamatergic markers are identified as excitatory and with enrichment of at least one of the GABA/glycinergic markers are identified as inhibitory. Clusters that only show enrichment for serotonergic, cholinergic, or a mix of glutamatergic and glycinergic markers are labelled as “Other”. The neuron cluster that does not express any of the previously mentioned marker genes is labelled “Unidentified”. **(C)** Pie charts show the proportions of excitatory, inhibitory, and “Other” neuron populations at 0, 1, 3 and 6 wpi. **(D)** *in silico* calculation of E/I ratios during SC regeneration. Bar charts depict the ratio between excitatory and inhibitory neurons at 0, 1, 3 and 6 wpi. Dots indicate single biological replicates. Two biological replicates are shown per time point. Error bars represent SEM. **(E, F)** Immunostaining for RFP, GFP, HuC/D and Hoechst in SC cross sections from Tg(*vglut2a*:RFP) and Tg(*gad1b*:GFP) zebrafish at 0, 1, 3 and 6 wpi. White arrows point to RFP^+^ HuC/D^+^ neurons in E and GFP^+^ HuC/D^+^ neurons in F. Insets depict single channel views of *vglut2a*:RFP or *gad1b*:GFP in dorsal and ventral SC domains, respectively. **(G, H)** Quantification of glutamatergic and GABAergic neurons after SCI. Percent *vglut2a*^+^ HuC/D^+^ neurons (ANOVA p-value: 0.0086) and *gad1b*^+^ HuC/D^+^ neurons (ANOVA p-value: 0.0043) were normalized to the total number of HuC/D^+^ neurons. **(I)** *in vivo* quantification of E/I ratios during SC regeneration. Ratios of glutamatergic to GABAergic neurons were calculated at 0, 1, 3 and 6 wpi (ANOVA p-value: 0.03421). Solid circles and polygons in bar charts indicate individual animals and sample sizes are indicated in parentheses. SC cross sections 450 µm rostral to the lesion were analyzed. Brown-Forsythe and Welch ANOVA test was performed in G, H and I with Dunnett’s T3 multiple comparisons test performed across different time points having 95% CI. * p≤0.05, ** p ≤0.01; Scale bars, 50 µm.

To validate our bioinformatic E/I balance calculations *in vivo*, we generated dual transgenic reporter fish that simultaneously label glutamatergic excitatory neurons (*vglut2a*:RFP) and GABAergic inhibitory neurons (*gad1b*:GFP) (Satou et al., 2013). Complete SC transections were performed on double transgenic animals and SC tissues were collected for histological examination at 0, 1, 3 and 6 wpi (Fig. 3E-F). Absolute numbers and relative proportions of RFP-, GFP- and HuC/D-expressing neurons were quantified in SC cross sections at 450 μm rostral to the lesion to evaluate the E/I landscape during regeneration. Compared to control SC tissues, the profiles of *vglut2a* ^+^ excitatory neurons (*vglut2a* ^+^ HuC/D^+^) doubled in proportion at 1 wpi and were maintained near 1 wpi levels until 6 wpi (Fig. 3G and S3F). On the other hand, the profiles of *gad1b*^+^ inhibitory neurons (*gad1b*^+^ HuC/D^+^) increased by 50% at 6 wpi relative to controls (Fig. 3H and S3G). Notably, the absolute numbers of HuC/D^+^ neurons drastically increased at 1 and 3 wpi, indicating extensive neuronal remodeling during SC regeneration (Fig. S3H). Thus, consistent with our snRNA-seq findings, *in vivo* studies confirmed an early surge in excitatory neurons at 1 wpi and a recovery of the E/I balance toward baseline levels by 6 wpi (Fig. 3I).

### Neurogenesis of excitatory and inhibitory neurons after spinal cord injury

We explored the cellular bases behind the recovery of E/I balance during zebrafish SC repair. The observed E/I ratio changes could be due to sequential neurogenesis of excitatory then inhibitory neurons after SCI. Alternatively, E/I ratios could reflect plasticity and changes in the neurotransmitter properties of pre-existing neurons. To distinguish between these possibilities, we first performed a cell proliferation assay. Dual transgenic fish for *vglut2a*:RFP and *gad1b*:GFP were subjected to SCI and a single EdU pulse for 24 hrs prior to SC tissue collection. SC tissues were harvested at 1, 3 and 6 wpi for co-labelling with the proliferation marker EdU (Fig. 4A-B and S4A). SC tissues from uninjured animals that received a single EdU injection for 24 hrs were harvested as controls. In striking manifestation of the regenerative capacity of zebrafish SC tissues after SCI, EdU incorporation was markedly elevated at 1 wpi relative to 0, 3 and 6 wpi (Fig. 4C and S4B). The profiles and absolute numbers of newly generated glutamatergic neurons (*vglut2a*^+^ EdU^+^) peaked at 1 wpi relative to control, 3 or 6 wpi SC sections (Fig. 4D and S4C), supporting a surge of regenerating excitatory neurons at 1 wpi. As the profiles of GABAergic neurons showed a gradual increase up to 6 wpi (Fig. 3H) and due to low numbers of EdU^+^ cells at 3 and 6 wpi, we were unable to evaluate the profiles of newly generated GABAergic neurons with a single EdU pulse (Fig. 4E and S4D).

**Figure 4.**
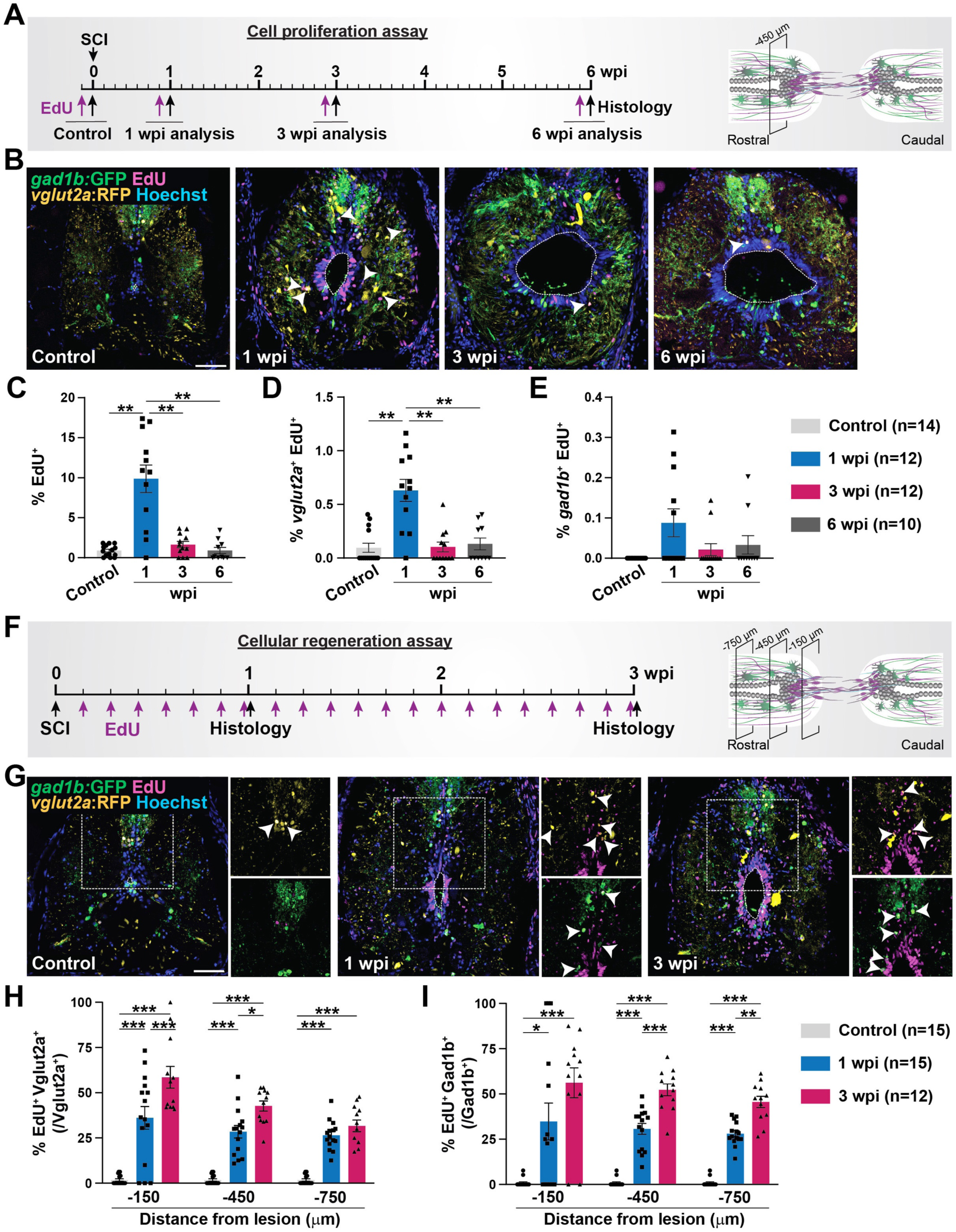
Regeneration of excitatory and inhibitory neurons after SCI. **(A)** Experimental timeline to assess cell proliferation at 1, 3 and 6 wpi. Tg(*vglut2a*:RFP; *gad1b*:GFP) zebrafish were subjected to SC transections. Single intraperitoneal EdU injections were performed at either 6, 20, or 41 days post-injury. SC tissues were harvested for analysis at 1, 3, or 6 wpi and at 24 hours after EdU injection. Uninjured animals were injected with EdU and collected 24 hours after EdU injection as controls. SC cross-section 450 µm rostral to the lesion was used for quantifications. **(B)** Staining for RFP, GFP, EdU and Hoechst in SC cross sections of Tg(*vglut2a*:RFP; *gad1b*:GFP) zebrafish at 0, 1, 3 and 6 wpi. White arrows indicate *vglut2a*^+^ EdU^+^ neurons. Dotted lines delineate central canal edges. **(C-E)** Quantification of EdU^+^ cells (ANOVA p-value <0.0001), *vglut2a*^+^ EdU^+^ neurons (ANOVA p-value <0.0001) and *gad1b*^+^ EdU^+^ neurons (ANOVA p-value: 0.0449) at 0, 1, 3 and 6 wpi. Percent cells were normalized to the total number of nuclei in SC sections. **(F)** Experimental timeline to assess the cumulative profiles of regenerating neurons at 1 and 3 wpi. Tg(*vglut2a*:RFP; *gad1b*:GFP) zebrafish were subjected to SC transections and daily intraperitoneal EdU injections. SC tissues were harvested for analysis at 1 and 3 wpi. Control SCs received daily EdU injection for 7 days before collection. Tissue sections 150, 450 and 750 μm rostral to the lesion were analyzed. **(G)** Staining for RFP, GFP, EdU and Hoechst in SC cross sections of Tg(*vglut2a*:RFP; *gad1b*:GFP) zebrafish at 0, 1 and 3 wpi. White arrows indicate *vglut2a*^+^ EdU^+^ neurons. Insets show single channel views of *vglut2a*:RFP and *gad1b*:GFP expression. Dotted lines delineate central canal edges. **(H, I)** Quantification of regenerating glutamatergic neurons (*vglut2a*^+^ EdU^+^) (ANOVA p-value <0.0001) and regenerating GABAergic neurons (*gad1b*^+^ EdU^+^) (ANOVA p-value <0.0001). For each section, percent cells were normalized to the number of *vglut2a*^+^ EdU^+^ neurons in H and *vglut2a*^+^ EdU^+^ neurons in I. Brown-Forsythe and Welch ANOVA test were performed in C, D and E with Dunnett’s T3 multiple comparisons test performed across different time points with 95% CI. Two-way ANOVA was performed in H and I with Tukey’s multiple comparison test having 95% CI. * p≤0.05, ** p ≤0.01, *** p≤0.001. Solid circles and polygons in the bar charts indicate individual animals and sample sizes are indicated in parentheses. Scale bars, 50 µm.

To overcome this limitation and evaluate the numbers of regenerating glutamatergic and GABAergic neurons after SCI, we performed SC transections on *vglut2a*:RFP and *gad1b*:GFP dual transgenic fish followed by daily EdU injections for 1 or 3 wpi (Fig. 4F-G and S4E). Uninjured control animals received daily EdU injections for 1 week prior to SC tissue collection. Compared to single EdU pulses, daily EdU labeling allowed us to estimate the total and cumulative profiles of regenerating neurons. To account for anatomical differences along the proximo-distal axis of lesioned SC tissues, RFP-, GFP- and EdU-expressing cells were quantified from SC sections 150, 450 and 750 μm rostral to the lesion. The numbers and proportions of EdU^+^ cells sharply increased at 1 wpi and showed less pronounced additive EdU incorporation at 3 wpi (Fig. S4F-G). The profiles of newly formed glutamatergic (EdU^+^ *vglut2a*^+^) neurons markedly increased at 1 and 3 wpi relative to controls (Fig. 4H and S4H). The profiles of regenerating GABAergic (EdU^+^ *gad1b*^+^) neurons were significantly elevated at 1 wpi compared to controls, and continued to increase at 3 wpi compared to 1 wpi (Fig. 4I and S4I). As proximal tissue sections (150 μm from lesion) showed elevated neurogenesis across time points, differential dynamics of glutamatergic and GABergic neuron regeneration were most prominent in more distal SC sections (750 μm from lesion) (Fig. 4H and 4I). At these levels, the profiles of newly formed glutamatergic (EdU^+^ *vglut2a*^+^) neurons sharply increased at 1 wpi and stabilized between 1 and 3 wpi (Fig. 4H), whereas GABAergic (EdU^+^ *gad1b*^+^) continuously increased from 1 to 3 wpi (Fig. 4I). These *in vivo* studies showed glutamatergic neurogenesis precedes a continuous and slower regeneration timeline for GABAergic neurons, suggesting that sequential neurogenesis of different neuron subtypes around the lesion accounts for the dynamic changes in E/I properties during zebrafish SC regeneration.

### Identification of injury-responsive iNeurons during early stages of spinal cord repair

To delve into the distribution of specific neuronal populations prior to and after SCI, we calculated the numbers and relative proportions of each neuron subcluster between 0 and 6 wpi (Fig. 5A, 3A and S3A-B). We also applied established marker genes for zebrafish larval neurons (Cavone *et al*., 2021) onto our marker scoring algorithm, identifying 7 major neuron subpopulations with varying cellular dynamics during regeneration (Fig. 5B, S5A-C and Table S8). Out of 30 neuronal clusters identified in our integrated dataset, cell proportions decreased >2-fold in 9 clusters (N0, 5, 10, 14, 15, 24, 25, 28 and 29), representing interneurons, V0, V2a interneurons, and motor neurons. On the other hand, 4 neuronal clusters expanded >2-fold after SCI (N18, 20, 23 and 27) (Fig. 5A). The numbers and proportions of N18 neurons increased 2.5-fold at 3 wpi and returned to control levels at 6 wpi (Fig. 5A). Consistent with delayed GABAergic neurogenesis (Fig. 3 and 4), Cluster N18 cells were identified as GABAergic V2b interneurons (Fig. 3B and 5B). The profiles of N20 neurons sharply increased 6.7-fold at 1 wpi compared to all other time points. Intriguingly, N20 was the only neuron cluster corresponding to neuroblast-like neurons, suggesting a pro-regenerative role for N20 neurons after SCI (Fig. 5A-B and S5A). Yet, the exact identities and role of N20 during SC repair are unknown. N23 neurons were injury-induced, accounting for 2% of total neurons at 1 wpi and decreasing to 0.6% of neurons at 6 wpi (Fig. 5A). N23 neurons were identified as motor neurons, which are known to regenerate via neurogenesis in adult zebrafish (Reimer *et al*., 2008). N27 neurons increased 2.9-fold at 1 wpi compared to other time points and were identified as serotonergic interneurons (Fig. 3B and 5B). We conclude that cluster N27 corresponds to intraspinal serotonergic neurons, which promote excitatory axon regrowth and coordinated locomotor recovery in zebrafish (Huang et al., 2021). This analysis revealed known and unanticipated neuronal dynamics during innate SC repair.

**Figure 5.**
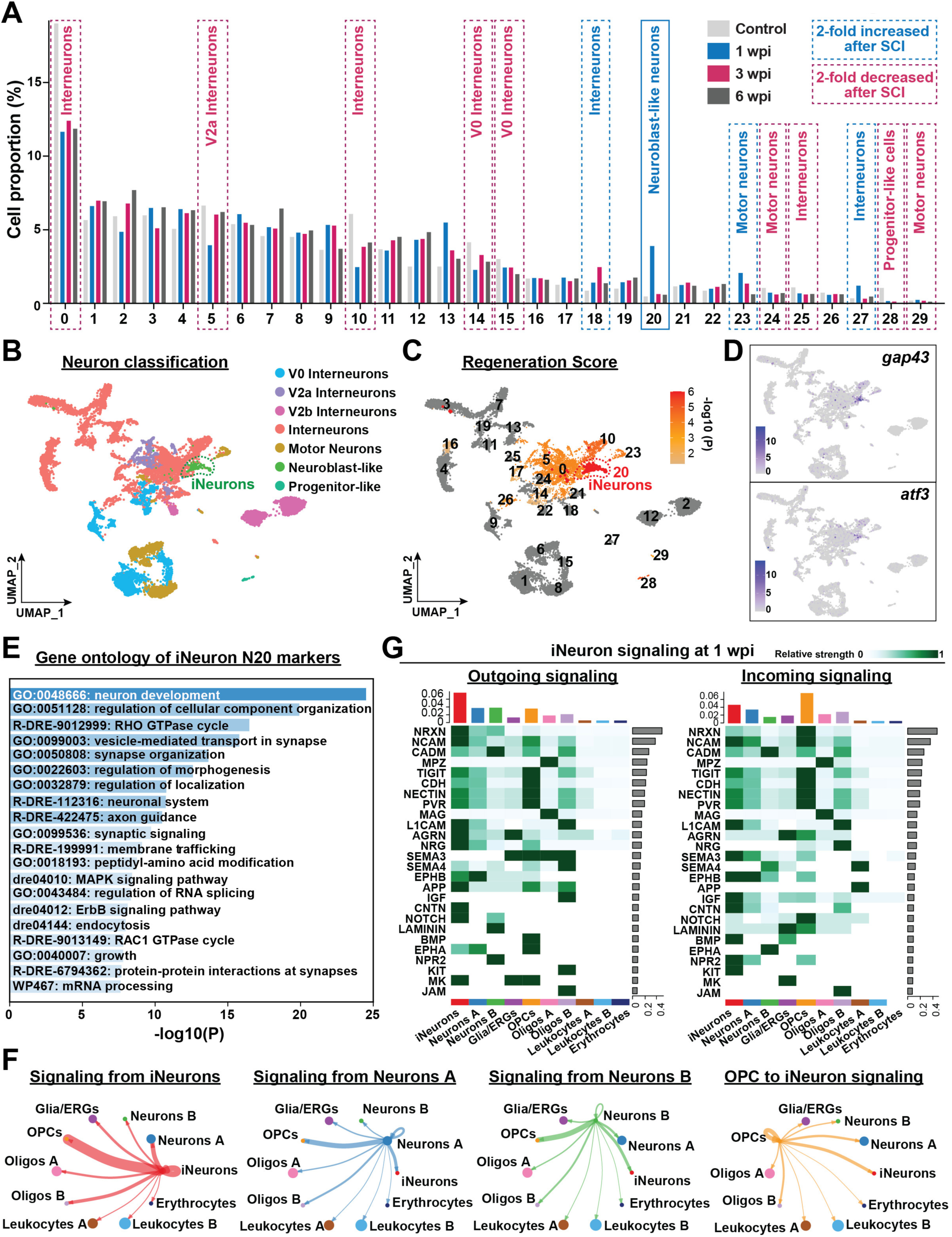
Emergence of an injury-responsive iNeuron signature during SC regeneration. **(A)** Dynamics of neuronal subclusters at 0, 1, 3 and 6 wpi. Cell proportions for each cluster and time point were normalized to the total number of neurons analyzed at that time point. Clusters that either increased or decreased by >2-fold in proportion are highlighted in red and blue, respectively. **(B)** UMAP plot showing predicted cluster identity for each neuron subpopulations. V0, V2a, V2b, interneurons, motor neurons are shown along with clusters identified as neuroblast-like or progenitor-like neurons. Cluster N20 was the only cluster identified as neuroblast-like neurons. **(C)** UMAP plot showing Regeneration score for each neuron subclusters. −log10P value of each cluster is represented in a gradient scale of gray (min=0) to red (max=6). Cluster N20 showed the highest regeneration score compared to all other neurons. **(D)** Feature plots show the expression of the regeneration associated genes *gap43* and *atf3* in cluster N20. **(E)** Gene ontology of N20 iNeuron DE markers. Twenty of the most enriched terms are shown. X-axis represents −log10(p-value) for each term. **(F)** Circle plots represent cell-cell interaction strengths at 1 wpi. Arrow thickness is directly proportional to the cumulative strengths of all predicted interactions between clusters. **(G)** Signaling pathways outgoing from or received by iNeurons at 1 wpi. Bar graphs at the top of each heatmap show cumulative signaling strengths per cell population. Bar graphs at the right of each heatmap show cumulative signaling strength per pathway.

To estimate the regenerative potential of spinal neurons after SCI, we assigned a “regeneration score” to individual neuron clusters by cross referencing the DE markers for each cluster with a compiled list of regeneration associated genes. RAGs were defined based on their association with the GO term “regeneration” in the Amigo database (http://amigo.geneontology.org/amigo/landing) (Tables S4 and S5). Strikingly, compared to all other neuron subtypes, cluster N20 showed the highest regeneration score (Fig. 5C and S5D) and expressed known RAG genes including *gap43* and *atf3* (Fig. 5D) (Curtis et al., 1993; Doster et al., 1991; Gey et al., 2016; Jankowski et al., 2009; Jing et al., 2012; Kole et al., 2020; Wang et al., 2017; Wang et al., 2015; Williams et al., 2015; Zhang et al., 2005). Gene ontology comparisons showed that terms associated with neuron development, synaptic organization and axon extension were enriched in N18, 20, 23 and 27 (Fig. S5E). However, regeneration associated gene enrichment was most pronounced and most significant for cluster N20 (Fig. S5E). We propose that cluster N20 represents a new, transient and unique population of injury-responsive neurons (iNeurons) that are likely to contribute to functional SC repair.

### iNeurons are signaling hubs during spinal cord regeneration

To evaluate the role of N20 iNeurons during SC regeneration, we determined the extent and identities of the signaling pathways that are initiated and received by iNeurons using CellChat. To specifically identify iNeurons (N20 cells) within the rest of the SC microenvironment, we first tracked and identified iNeurons in the integrated dataset that includes all SC cells (Fig. S5D). Cells from clusters 5 and 4 from the integrated dataset accounted for the majority of iNeurons and were renamed iNeurons for CellChat analysis. Strikingly, compared to the global cell-interactions derived from the integrated dataset, the cumulative and reciprocal signaling strengths were highest between iNeurons and OPCs at 1 wpi (Fig. 5F). The cumulative strengths of outgoing and incoming signaling were markedly lower in the rest of the neuronal cells compared to iNeurons (Fig. 5F). These findings suggest that iNeuron signaling accounts for the majority of neuronal signaling at 1 wpi.

To examine the molecular features of the crosstalk between iNeurons and OPCs, we identified major signaling networks that are either incoming or outgoing from iNeurons (Fig. 5G and S5F-G). CellChat analysis showed iNeurons have a high probability to transduce NCAM1/2, and NRXN3 signaling to OPCs and other neurons. TIGIT, NRXN1/2, CDH4 and CADM1/3 were also predicted as incoming and outgoing iNeuron signals. Consistent with the ontology of iNeuron expressed genes, we observed an enrichment for reciprocal Neuregulin (NRG)-ERBB signaling from iNeurons to OPCs. ErbB signaling was recently shown to regulate OPC transformation and spontaneous remyelination after SCI in mice (Bartus et al., 2019). We found NRG1 and NRG2 ligands activate of the ERBB3/4 receptor mediated signaling between iNeurons and OPCs (Fig. 6D). These results showed iNeurons are potent centers of molecular signaling during SC regeneration and suggested that reciprocal signaling between iNeurons and OPCs is a principal feature of innate SC repair.

**Figure 6.**
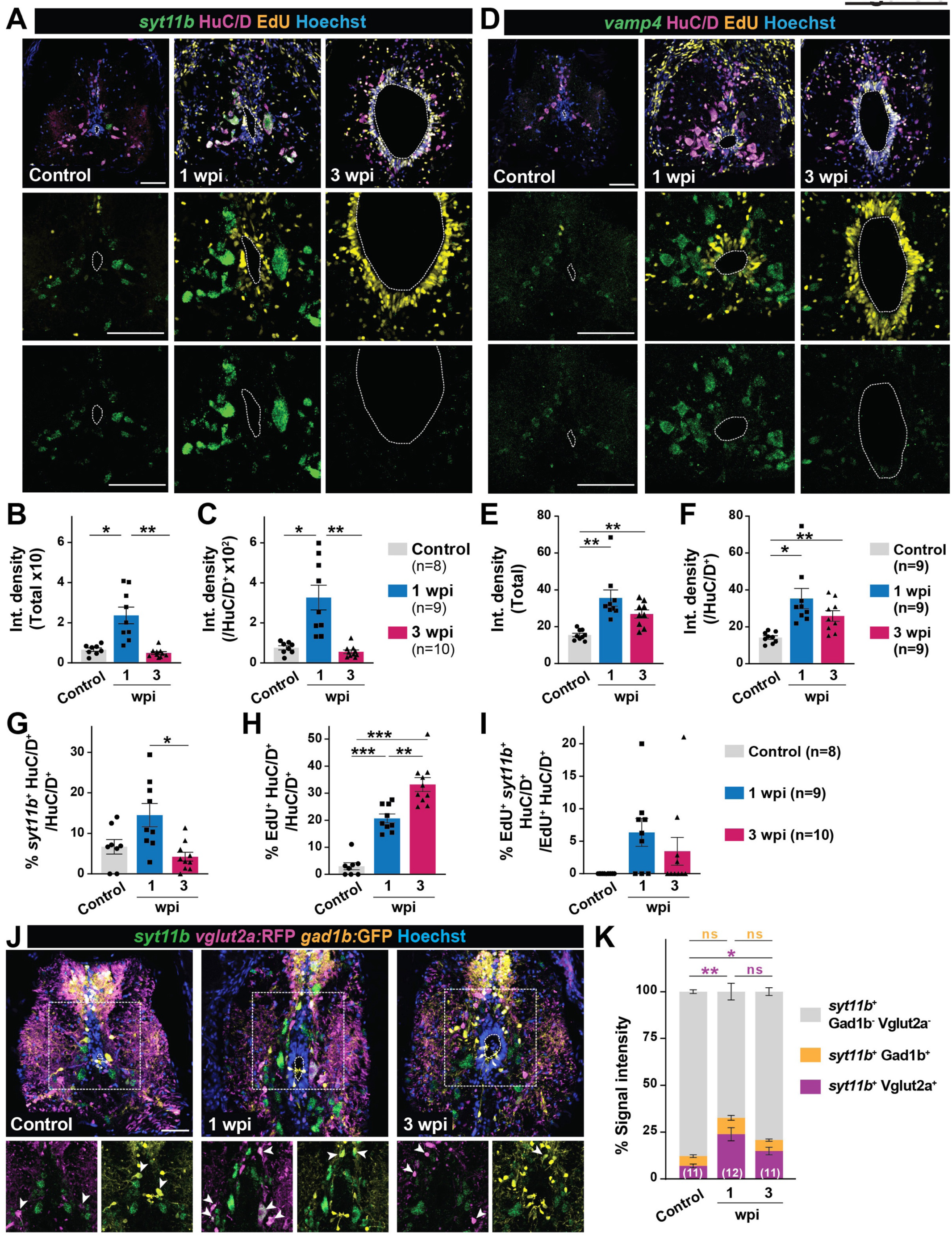
*in vivo* mapping of iNeurons in zebrafish spinal tissues. **(A)** HCR *in situ* hybridization for *syt11b* and staining for HuC/D, Hoechst and EdU were performed on wild-type SC cross sections at 0, 1 and 3 wpi. Dotted lines delineate central canal edges. **(B, C)** Quantification of *syt11b* HCR *in situ* hybridization. Integrated density was calculated in the whole SC cross sections (ANOVA p-value: 0.0007) and inside HuC/D^+^ neurons (ANOVA p-value: 0.0009) at 0, 1 and 3 wpi. **(D)** HCR *in situ* hybridization for *vamp4* and staining for HuC/D, Hoechst and EdU were performed on wild-type SC cross sections at 0, 1 and 3 wpi. Dotted lines delineate central canal edges. **(E, F)** Quantification of *vamp4* HCR *in situ* hybridization. Integrated density was calculated in the whole SC cross sections (ANOVA p-value: 0.0012) and inside HuC/D^+^ neurons (ANOVA p-value: 0.0043) at 0, 1 and 3 wpi. **(G-I)** Quantification of *syt11b* expressing cells after SCI. Percent *syt11b*^+^ HuC/D^+^ neurons (normalized to HuC/D^+^ neurons in G) (ANOVA p-value: 0.0062), percent EdU^+^ HuC/D^+^ neurons (normalized to HuC/D^+^ neurons in H) (ANOVA p-value <0.0001), and percent *syt11b*^+^ EdU^+^ HuC/D^+^ neurons (normalized to EdU^+^ HuC/D^+^ neurons in I) (ANOVA p-value: 0.0813). Analysis was performed on SC cross sections at 0, 1 and 3 wpi. **(J)** HCR *in situ* hybridization for *syt11b* in SC cross-sections of Tg(*vglut2a*:RFP; *gad1b*:GFP) zebrafish at 0, 1 and 3 wpi. Dotted lines delineate central canal edges. Arrowheads point to overlapping of *syt11b* transcripts with either *vglut2a* or *gad1b*. **(K)** Quantification of *syt11b* transcripts within *vglut2a*^+^ and *gad1b*^+^ neurons at 0, 1 and 3 wpi (ANOVA p-value <0.0001). *syt11b* signal outside either of these neurotransmitter populations was also quantified. SC cross sections at 450 µm rostral to the lesion were analyzed. Solid circles and polygons in the bar charts indicate individual animals and sample sizes are indicated in parentheses. Brown-Forsythe and Welch ANOVA test was performed in B, C, E, F, G, H, and I with Dunnett’s T3 multiple comparisons test performed across different time points with 95% CI. Two-way ANOVA was performed in K with Tukey’s multiple comparisons test with 95 % CI. * p≤0.05, ** p ≤0.01, *** p≤0.001. Scale bars, 50 µm.

### *in vivo* mapping of iNeurons during SC regeneration

To validate the emergence of an iNeuron transcriptional signature in regenerating SC tissues, we performed HCR *in situ* hybridization for select markers that are uniquely expressed in iNeurons compared to all other spinal cell types in the integrated dataset (Fig. S6A). To enable differential labeling of regenerating and pre-existing neurons, wild-type fish were subjected to SCI and daily EdU injections. *syt11b* and *vamp4* HCR *in situ* hybridization and staining for HuC/D and EdU were performed at 0, 1 and 3 wpi (Fig. 6A and 6D). To test whether iNeuron markers are preferentially expressed in neurons, integrated densities of *in situ* hybridization signals were quantified in total SC tissues and in HuC/D^+^ neurons (Fig. 6A-F and S6B-H). At 1 wpi, *synaptotagmin XI b (syt11b)* expression increased 3.6-fold in total SC tissues and 4.3-fold in HuC/D^+^ neurons (Fig. 6B-C). *syt11b* expression was reduced to wild-type levels by 3 wpi (Fig. 6B-C). *vesicle-associated membrane protein 4 (vamp4)* transcripts were upregulated at 1 and 3 wpi, with 2.3- to 2.5-fold increase in total SC tissues and HuC/D^+^ neurons at 1 wpi (Fig. 6E-F). Since *syt11b* showed robust expression and upregulation in neurons and to address putative redundancy between the *syt11b* and *syt11a* paralogs, we examined *syt11a* expression after SCI (Fig. S6E-H). Like *syt11b*, *syt11a* transcripts were upregulated after injury, including in neurons. However, unlike *syt11b*, *syt11a* expression was not restricted to neurons including expression in ependymal cells surrounding the central canal at 1 wpi (Fig. S6E). These findings validated *syt11b* and *vamp4* as markers of zebrafish iNeurons during SC regeneration.

### iNeurons are surviving neurons that elicit spontaneous plasticity after injury

The emergence of an iNeuron transcriptional profile at 1 wpi raised a central question related to their cell of origin and pro-regenerative signature. We postulated that iNeurons could either represent newly formed regenerating neurons, or a population of spontaneously plastic neurons that survive injury and support immediate repair. To distinguish between these possibilities, we quantified the profiles of iNeurons that express *syt11b/a*, HuC/D and EdU (Fig. 6G-I and S6F-H). In these experiments, HuC/D^+^ EdU^+^ represent regenerating neurons and HuC/D^+^ EdU^-^ represent pre-existing neurons. Expectedly, the profiles of regenerating neurons significantly increased from 1 to 3 wpi, indicating potent neurogenic responses are activated during innate SC repair (Fig. 6H). At 1 wpi, *syt11b* transcripts labeled 14.5% of HuC/D^+^ neurons (Fig. 6G), and only 6.4% of regenerating neurons (Fig. 6I). These results indicated that over 90% of iNeurons are pre-existing neurons that survive SCI.

Since the majority of iNeurons are pre-existing in control SC tissues, we examined whether iNeurons represent a specific population of neurons with elevated survival capacity or spontaneous plasticity. To address this question, we performed *syt11b* HCR *in situ* hybridization on *vglut2a*:RFP and *gad1b*:GFP dual transgenic fish, which simultaneously label glutamatergic and GABAergic neurons (Fig. 6J). Neuron subtypes not labeled with *vglut2a*:RFP or *gad1b*:GFP were labeled as others. The integrated density of *syt11b* transcript signals broadly increased at 1 wpi (Fig. S6I), including upregulated expression in *vglut2a* ^+^ and *gad1b*^+^ neurons at 1 wpi (Fig. S6J-K). Notably, only 37.5% and 20.7% of the combined *syt11b* signal co-localized to glutamatergic and GABAergic neurons at 1 and 2 wpi, indicating that “other” neuron subtypes express the majority of *syt11b* during regeneration (Fig. 6K). We also performed *syt11b* HCR *in situ* hybridization along with Hb9, *isl1:GFP* or Pax2 staining (Fig. S6L), and found that 30-40% of Hb9^+^, *isl1:GFP^+^*and Pax2^+^ neurons expressed *syt11b* at 1wpi (Fig. S6M). These findings indicated that iNeurons do not represent a specific neuron subtype and suggested that multiple zebrafish neurons survive SCI and elicit an iNeuron transcriptional and regenerative signature during early stages of SC repair.

### iNeurons are integrated in the spinal circuitry after spinal cord injury

We performed two sets of tracing experiments to trace axons projecting across the lesion using either rostral (Fig. 7A-C) or caudal labeling (Fig. S7A-C) with Biocytin at 1 wpi. For rostral labeling, a Biocytin-soaked gelfoam was applied at the brainstem level at 1 wpi and 3 hours prior to collecting SC tissues. In addition to Biocytin, *syt11b* HCR *in situ* hybridization and HuC/D staining were performed for simultaneous labeling of iNeurons and total neurons, respectively (Fig. 7A). We found that 28-33% of HuC/D^+^ neurons between 450 and 750 μm caudal to the lesion were labeled (Fig. 7B), indicating integration of caudal neurons into the spinal circuitry as early as 1 wpi. 20-38% of rostrally-labeled caudal neurons (Biocytin^+^ HuC/D^+^) expressed *syt11b* (Biocytin^+^ HuC/D^+^ *syt11b*^+^) (Fig. 7C), suggesting *syt11b*^+^ iNeurons are part of the integrated SC circuitry at 1wpi. To validate these results, we also applied a Biocytin-soaked gelfoam caudal to the lesion at 1 wpi and collected SC tissues for histology 3 hours post-labeling (Fig. S7A). Caudal tracing labeled 18-20% of HuC/D^+^ neurons 450 and 750 μm rostral to the lesion (Biocytin^+^ HuC/D^+^) (Fig. S7B), with ∼40% of these neurons expressing *syt11b* (Biocytin^+^ HuC/D^+^ *syt11b*^+^) (Fig. S7C). These studies confirmed that iNeurons establish functional connections into the regenerating SC circuitry as early as 1 wpi.

**Figure 7.**
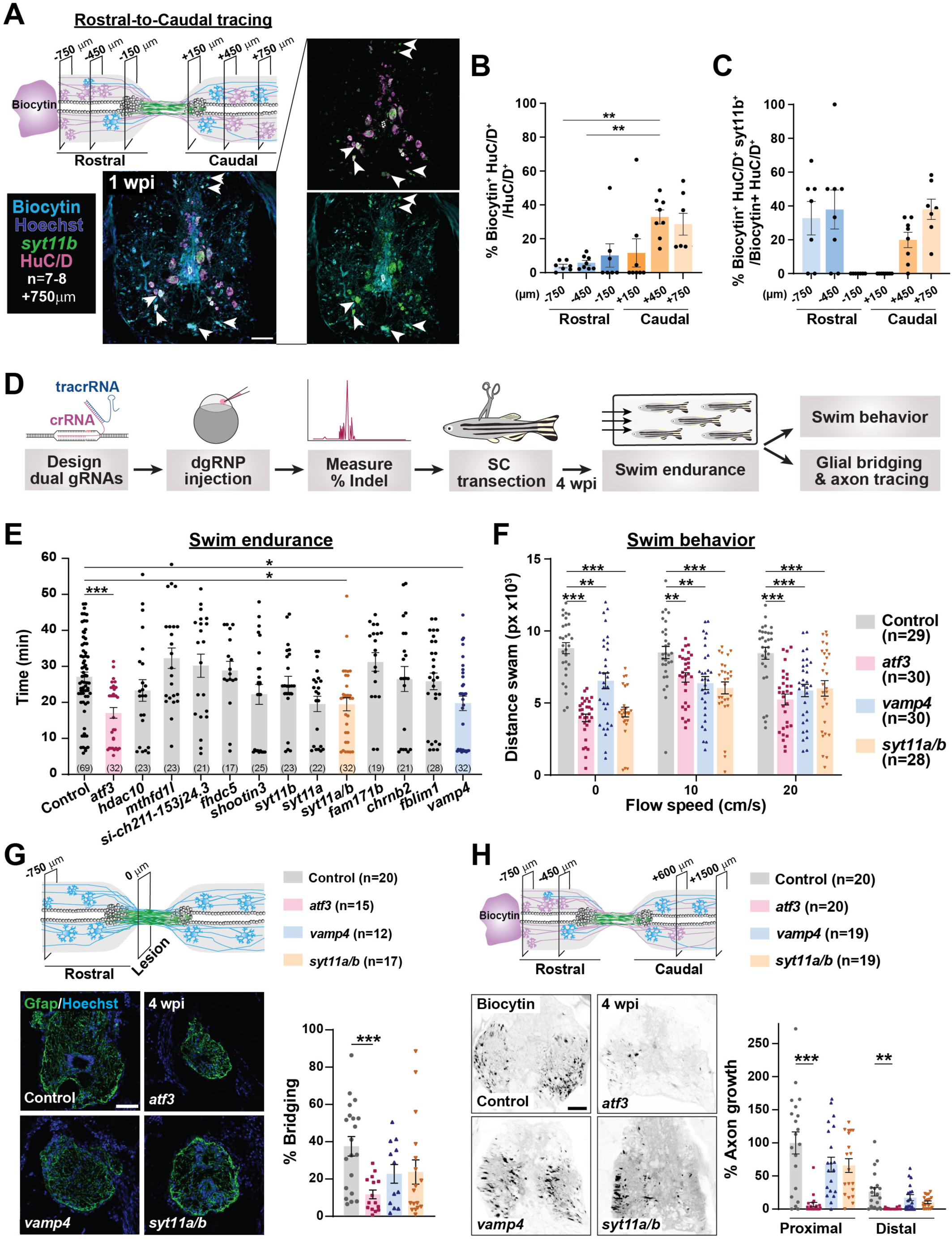
Tracing and CRISPR/Cas9 mutagenesis of iNeuron marker genes in zebrafish. **(A-C)** Rostral-to-caudal tracing of neurons at 1 wpi. Biocytin was applied 4 mm rostral to the lesion and co-labeled with *syt11b* HCR *in situ* hybridization and HuC/D staining. SC cross sections 750 mm caudal to the lesion are shown in A. Dotted lines delineate central canal edges. The profiles of Biocytin-labeled neurons (Biocytin^+^ HuC/D^+^) were normalized to total neurons (HuC/D^+^) in B. The profiles of Biocytin-labeled neurons (Biocytin^+^ HuC/D^+^) were normalized to total neurons (HuC/D^+^) in B. The profiles of Biocytin-labeled iNeurons (Biocytin^+^ HuC/D^+^ *syt11b*^+^) were normalized to traced neurons (Biocytin^+^ HuC/D^+^) in C. ANOVA p-values are 0.0029 for A and 0.0011 for B. **(D)** Pipeline for CRISPR/Cas9 mutagenesis of iNeuron marker genes for regeneration assessment. CRISPR/Cas9 dual-guide ribonucleic protein duplexes targeting iNeuron genes injected into one-cell zebrafish embryos. Mutagenesis efficiency was confirmed by capillary electrophoresis and animals with >90% mutagenesis were used for SCI. Swim endurance, swim behavior and histological regeneration assays were performed at 4 wpi. **(E)** Functional recovery in CRISPR/Cas9 targeted animals 4 wpi. For each group of targeted animals, uninjected siblings were subjected to SCI and swim endurance assays (ANOVA p-value <0.0001). Dots represent individual animals. **(F)** Swim behavior assays in CRISPR/Cas9 targeted animals 4 wpi (ANOVA p-value <0.0001). Swim distance was tracked under water current velocities of 0, 10 and 20 cm/s. **(G)** Glial bridging in CRISPR/Cas9 targeted animals at 4 wpi. Representative immunohistochemistry shows the Gfap^+^ bridge at the lesion site in *atf3, vamp4* and *syt11a/b* crispants relative to controls. Percent bridging represents the cross-sectional area of the glial bridge at the lesion site (0 µm) relative to the intact SC 750 µm rostral to the lesion (ANOVA p-value: 0.0049). **(H)** Anterograde axon tracing in CRISPR/Cas9 targeted animals at 4 wpi. Biocytin axon tracer was applied rostrally and analyzed at 600 μm (proximal) (ANOVA p-value <0.0001) and 1500 μm (distal) (ANOVA p-value: 0.0006) caudal to the lesion. Representative traces of biocytin are shown at the proximal level. Axon growth was first normalized to the extent of biocytin at 450 and 750 µm rostral to the lesion, and then to labeling in wild-type controls at the proximal level. Solid circles and polygons in the bar charts indicate individual animal and sample sizes are indicated in parentheses. Brown-Forsythe and Welch ANOVA test was performed in B, E and G with Dunnett’s T3 multiple comparisons test performed across different time points with 95% CI. Two-way ANOVA with Dunnett’s multiple comparisons test (95% CI) was performed in C. * p≤0.05, ** p ≤0.01, *** p≤0.001. Scale bars, 50 µm.

### iNeuron genes are required for cellular and functional recovery after spinal cord injury

To evaluate the roles of iNeuron marker genes during SC regeneration, we employed CRISPR/Cas9 mutagenesis, SCI and functional swim assays (Fig. 7D) (Burris *et al*., 2021; Klatt Shaw and Mokalled, 2021). Among the top DE markers of N20 iNeurons, 10 genes were filtered as enriched in iNeurons and chosen for mutagenesis (*hdac10, mthfd1l, si:ch211-153j24.3, fhdc5, shootin3, syt11b, fam171b, chrnb2b, fblim1, vamp4*) (Fig. S7D). *atf3* was selected as a positive control (Wang *et al*., 2017). In addition to targeting *syt11b*, we individually and combinatorially targeted the *syt11a* and *syt11b* paralogs to account for putatively redundant effects on regeneration. For efficient and simultaneous gene targeting in zebrafish, we used a recently adapted CRISPR/Cas9 protocol that achieves near complete mutagenesis in F0 injected adults (Burris *et al*., 2021; Klatt Shaw and Mokalled, 2021). CRISPR/Cas9 targeted animals (crispants) were raised to adulthood and mutagenesis rates were examined by capillary electrophoresis (Fig. S7E). Adult crispants with >90% mutagenesis were subjected to SCI, and swam against increasing water current inside an enclosed swim tunnel to screen for functional regeneration defects. At 4 wpi, swim endurance was significantly reduced in *atf3* and *vamp4* crispants (Fig. 7E). Targeting *syt11a and syt11b* paralogs individually did not alter swim endurance. However, swim endurance was diminished in *syt11a;syt11b* dual crispants, indicating the *syt11* paralogs were functionally redundant during SC repair (Fig. 7E). Importantly, overall growth and behavior were comparable between crispant fish and their control siblings prior to SCI, indicating the swim defects observed in these crispants and injury-induced. To validate swim capacity defects in iNeuron gene crispants, we tracked the swim behavior of *atf3*, *vamp4* and *syt11a;syt11b* crispants under at 0, 10 and 20 10 cm/sec current velocities (Fig. 7F). Compared to uninjected control siblings and in the absence of water flow, swim distance was 50-55% reduced in *atf3* and *syt11a;syt11b* crispants and 25% reduced in *vamp4* crispants (Fig. 7F). Swim distance was significantly lower in *atf3*, *vamp4* and *syt11a/b* at 10 and 20 cm/sec water current velocities (Fig. 7F). These results indicated the iNeuron markers *atf3*, *vamp4* and *syt11a/b* are required for functional SC repair in zebrafish.

To date, cellular growth across the lesion has served as a primary readout of cellular regeneration in zebrafish. Glial bridging and axon tracing assays were performed to evaluate the extents of glial and axonal regeneration in iNeuron gene crispants. By Gfap immunostaining, the cross-sectional area of glial bridges was 65% reduced in *atf3* cripants at 4 wpi, but was not significantly affected in *vamp4* and *syt11a;syt11b* crispants compared to controls (Fig. 7G). At this time point, anterograde axon tracing using Biocytin showed comparable axon regrowth from hindbrain neurons into the proximal and distal SC sections between *vamp4*, *syt11a;syt11b* and control animals (Fig. 7H). Consistent with their defective glial bridging, axon regrowth into the proximal and distal sections of caudal SC tissues was markedly impaired in *atf3* cripants. Thus, *vamp4* and *syt11a/b* are required for functional SC repair but dispensable for cellular regeneration across the lesion.

These studies showed iNeurons play essential regeneration-independent plasticity-based roles after SCI, and indicated vesicular trafficking is required for the recovery of local neuronal circuitry during innate SC repair.

## DISCUSSION

This study presents a single-cell atlas of innate SC repair in adult zebrafish, uncovers neurons as potent signaling hubs after SCI, identifies cooperative modes of regeneration- and plasticity-based neuronal repair, and shows essential roles for vesicular trafficking in spontaneous neuronal plasticity.

The single-cell dataset generated in this study offers a comprehensive resource to determine regenerative cell identities and mechanisms in highly regenerative vertebrates. The use of snRNA-seq is ideal to profile the majority of spinal cell types in zebrafish. Single-cell RNA sequencing is inherently biased against larger cell types such as neurons and astrocytes (Bakken et al., 2018; Lake et al., 2017; Wu et al., 2019). snRNA-seq reduces the challenges of cell dissociation, retrieving large numbers of neurons and glia. While dissection- and dissociation-induced gene expression changes in control samples present a longstanding difficulty for tissue regeneration studies, our protocol minimizes background activation of injury-induced genes by flash-freezing SC tissues from individual animals during tissue collection. Finally, large numbers of nuclei were obtained by pooling frozen SC tissues, minimizing confounding problems associated with snRNA-seq and allowing us to identify small cell populations (Bakken *et al*., 2018; Ding et al., 2020; Lacar et al., 2016; Lake *et al*., 2017; van den Brink et al., 2017). Overall, this dataset faithfully represents the proportions and dynamics of major cell populations during zebrafish SC regeneration.

Candidate approaches have unveiled central roles for several signaling pathways during SC regeneration (Cavone *et al*., 2021; Dias et al., 2012; Goldshmit *et al*., 2012; Reimer et al., 2009; Zhang et al., 2018). Yet, an integrated assessment of regenerative signaling pathways and global assessment of cell-cell interactions were missing. Using CellChat analysis, our single-cell dataset identified a myriad of cell-specific and injury-induced signaling pathways (Jin *et al*., 2021). Notably, comprehensive assessment of cell-cell signaling networks uncovered neurons as potent sources of signaling in homeostatic and lesioned SC tissues. Injury-induced neuronal signaling pathways include developmental pathways like FGF and WNT, extracellular matrix components like FN1, and axon guidance molecules like SEMA. Concerted activation of these injury-induced pathways is instrumental for neuron survival, neurogenesis, axon regrowth, proper axon guidance and synapse targeting (Briona *et al*., 2015; Garcia *et al*., 2018; Goldshmit *et al*., 2012; He *et al*., 2018; Saraswathy *et al*., 2022; Tome *et al*., 2023; Wehner et al., 2017). Further studies of injury-induced signaling pathways will shed light into signaling pathways that direct regeneration- or plasticity-driven SC repair.

SC tissues in adult zebrafish retain potent progenitor cells with radial glial features and established contribution to neurogenesis after SCI (Briona *et al*., 2015; Reimer *et al*., 2008; Saraswathy *et al*., 2022). SC regeneration is marked by early imbalance and late restoration of baseline E/I activity. Our data support a model in which excitatory neurons expand and repopulate regenerate SC tissues, while slower and continued neurogenesis of inhibitory neurons underlie the recovery of E/I balance at 6 wpi. These findings are consistent with published studies using a two-cut SCI model in adult zebrafish (Huang *et al*., 2021), and with the observation that *gad1b*:GFP and *vglut2a*:RFP expression domains are mutually exclusive in our SC sections. Genetic lineage tracing showed excitatory to inhibitory neurotransmitter switching has adverse effects on functional recovery after adult mouse SCI (Bertels *et al*., 2022). In zebrafish, fast motor neurons were shown to upregulate glutamate expression after injury or exercise (Bertuzzi et al., 2018). Although we cannot rule out similar mechanisms taking place during regeneration, we found that staggered increase in the profiles of newly generated excitatory and inhibitory neurons account for the recovery of E/I balance after SCI. We propose that in depth molecular evaluation of spinal progenitors is needed to better understand the potency and neurogenic capacities of neuronal progenitors during SC regeneration.

While zebrafish is an established model of neuron regeneration, our study establishes zebrafish as a renewed model to understand and manipulate fundamental mechanisms of plasticity-based neuronal repair. Neurogenesis-based neural repair is thought to be unattainable in mammals. Instead, the mammalian SCI field has directed its efforts to develop plasticity-based repair strategies. We propose that pre-existing zebrafish neurons upregulate an iNeuron transcriptional signature that includes regeneration associated genes, and that iNeurons elicit spontaneous plasticity during early stages of SC repair. The emergence of iNeurons after SCI is consistent with a recently identified population of embryonically derived dormant neurons that immediately respond to SCI in larval and adult zebrafish (Vandestadt *et al*., 2021). Similarity to the dormant precursor neurons reported by Vandestadt *et al.,* more than 90% of iNeurons are not newly generated. At 1 wpi, iNeuron signaling accounts for the majority of total neuronal signaling, eliciting particularly strong interactions with OPCs. We propose that future zebrafish studies will contribute insights and applications into mechanisms of plasticity-driven neural repair.

Unbiased characterization of the molecular players that direct iNeuron functions after SCI showed multiple components of vesicular trafficking are required for functional SC repair in zebrafish. Syt11a/b (synaptotagmin XI a/b) and Vamp4 (vesicle-associate membrane protein 4) are vesicle-associated proteins that play different roles in neuronal synapses. Syt11 is a non-canonical SNARE that inhibits spontaneous neurotransmission and bulk endocytosis, whereas Vamp4 promotes Calcium-dependent excitatory neurotransmission release and bulk endocytosis (Bakr et al., 2021; Li et al., 2021; Lin et al., 2020; Nicholson-Fish et al., 2015; Shimojo et al., 2019; Wang et al., 2016). Our CRISPR/Cas9 studies showed *syt11a/b* and *vamp4* are required for the recovery of swim function after SCI in zebrafish. These findings are consistent with recent mammalian studies showing that inhibition of the presynaptic release machinery is important for axon regrowth (Hilton et al., 2022), and that *Syt11* knockout in excitatory forebrain neurons impairs synaptic plasticity and memory (Shimojo *et al*., 2019). Our data reveals vesicular trafficking is an essential mechanism that underlies spontaneous plasticity and rapid neurite outgrowth in iNeurons. We propose that zebrafish provide a valuable model to investigate the origin of iNeurons, elucidate the role of vesicular trafficking in spontaneous plasticity, and identify new manipulations to promote plasticity after SCI.

## ACKNOWLEDGMENTS

We thank S. Ackerman, V. Cavalli and A. Johnson for discussion and the Washington University Zebrafish Shared Resource for animal care. We thank S. Nandagopal and T. Tsai for sharing the HCR probe design script. This research was supported by grants from the NIH (R01 NS113915 to M.H.M.) and a Postdoctoral Fellow Seed of Independence Grant from the Department of Developmental Biology at Washington University School of Medicine (to V.M.S.).

## ETHICS DECLARATIONS

The authors declare no competing interests.

## MATERIAL AND METHODS

### Zebrafish

Adult zebrafish of the Ekkwill, Tubingen and AB strains were maintained at the Washington University Zebrafish Core Facility. All animal experiments were performed in compliance with institutional animal protocols. Size matched (∼2 cm) male and female animals of ∼4 months of age were used. SC transection surgeries and regeneration analyses were performed in a blinded manner. For all experiments, 2 to 4 independent experimental replicates were performed using different clutches of animals. Within each experiment, experimental fish and control siblings of similar size and equal sex distribution were used. Transected animals from control and experimental groups were housed in equal numbers (4-7 fish) in 1.1-liter tanks. The following previously published zebrafish trains were used: Tg(*vglut2a*:RFP) and Tg(*gad1b*:GFP) (Satou *et al*., 2013).

### Spinal cord transection

Complete SC transections were performed as previously described (Mokalled *et al*., 2016). Briefly, zebrafish were anaesthetized in 0.2 g/L of MS-222 buffered to pH 7.0. Fine scissors were used to make a small incision that transects the SC 4 mm caudal to the brainstem region. Complete transection of the SC was visually confirmed at the time of surgery. Injured animals were also assessed at 2 or 3 dpi to confirm loss of swim capacity post-surgery.

### Isolation of spinal cord nuclei from adult zebrafish

For nuclear isolation, a previously described method to isolate nuclei from mouse SC tissues was adapted to zebrafish SCs (Matson *et al*., 2018). Three mm SC tissue, flanking the lesion site were collected from 50 adult zebrafish at time points 1, 3 and 6 wpi. Corresponding segments of 3 mm SC tissue were collected from uninjured control animals. For tissue lysis, the detergent mechanical lysis protocol described by Matson et al. 2018 was performed. SC tissues were homogenized at low setting for 15 seconds. Density gradient separation using sucrose solution was used to sediment the nuclei from the supernatant. Final nuclear lysates were resuspended using 100 µl of resuspension solution (1x PBS + 2% BSA + 0.2U/µl RNase inhibitor - New England Biolabs, Cat#: M0314S). Hoechst staining was performed to assess the quality of isolated nuclei based on their shape. Samples in which more than 70% of the nuclei were scored as ‘healthy’ and submitted for single nuclear RNA sequencing.

### Single nuclear RNA sequencing

For snRNA-seq, 30 µl of resuspension solution containing isolated nuclei at a concentration of ∼1000 nuclei/µl was submitted to Genome Technology Access Center at McDonnel Genome Institute of Washington University. Two biological replicates of each time points at 0, 1, 3 and 6 wpi were used. cDNA was prepared after the GEM generation and barcoding, followed by the GEM-RT reaction and bead cleanup steps. cDNA was amplified for 11-13 cycles then purified using SPRIselect beads. Purified cDNA samples were then run on a Bioanalyzer to determine the cDNA concentration. GEX libraries were prepared as recommended by the 10x Genomics Chromium Single Cell 3’ Reagent Kits User Guide (v3.1 Chemistry Dual Index) with appropriate modifications to the PCR cycles based on the calculated cDNA concentration. For sample preparation on the 10x Genomics platform, the Chromium Next GEM Single Cell 3’ Kit v3.1, 16 rxns (PN-1000268), Chromium Next GEM Chip G Single Cell Kit, 48 rxns (PN-1000120), and Dual Index Kit TT Set A, 96 rxns (PN-1000215) were used. The concentration of each library was accurately determined through qPCR utilizing the KAPA library Quantification Kit according to the manufacturer’s protocol (KAPA Biosystems/Roche) to produce cluster counts appropriate for the Illumina NovaSeq6000 instrument. Normalized libraries were sequenced on a NovaSeq6000 S4 Flow Cell using the XP workflow and a 50×10×16×150 sequencing recipe according to manufacturer protocol. A median sequencing depth of 50,000 reads/cell was targeted for each Gene Expression Library.

### Aligning snRNA-seq reads

After sequencing, the Illumina output was processed using the CellRanger (v6.0.0) recommended pipeline to generate gene-barcode count matrices. A custom reference genome was made with the “cellranger mkref” command, using the fasta file of zebrafish reference genome GRCz11 constructed from the Ensemble genome build (https://useast.ensembl.org/Danio_rerio/Info/Index) and the sorted Gene Transfer Format file (v4.3.2) from the improved zebrafish transcriptome annotation (Lawson *et al*., 2020). Base call files for each sample from Illumina were demultiplexed into FASTQ reads. Then, the “cellranger count” pipeline was used to align sequencing reads in FASTQ files to the custom reference genome. Both exon and intron sequences were included during the alignment. The filtered gene-barcode count matrices generated by “cellranger count” was used for downstream analysis.

### Quality Control

DecontX package (Yang *et al*., 2020) (celda v1.16.1) was used to remove droplets containing aberrant counts of ambient mRNA. The function “decontx” was used on the raw “RNA” counts to obtain “decontX” count data. Contamination score was assigned to each cell after running “decontx”. Cells with contamination score greater than 0.75 were filtered out and only “clean” cells (contamination score less than 0.75) were included for downstream analysis. Furthermore, decontX counts were used as default for pre-processing and normalization. After ambient RNA removal, DoubletFinder (McGinnis *et al*., 2019) (v2.0.3) was used to identify doublets formed from transcriptionally distinct cells. Optimal pK values were calculated from the outputs of function “paramsweep_v3”. Then, “doubletFinder_v3” function was used to predict doublet cells, where 50 principal components and pN value of 0.25 were given as input along with the previously calculated pK values (0.26). All the cells predicted as doublets were removed.

### Integrated analysis of snRNA-seq dataset

All the datasets were integrated and analyzed using Seurat (v4.1.1) package with R (v4.2.1) (R Core Team, 2018; Stuart et al., 2019). Each sample count matrix was filtered for genes that were expressed in at least 3 cells and cells expressing at least 200 genes, followed by cell quality assessment using commonly used QC matrixes (Ilicic et al., 2016). Cells having a unique number of genes between 200 to 4000 and a mitochondrial gene percentage <5 were used for downstream processing. Each dataset was independently normalized and scaled using the “SCTransform” function, which is an improved method for normalization, that performs a variance-stabilizing transformation using negative binomial regression (Hafemeister and Satija, 2019). Standard integration workflow of Seurat was used to identify shared sources of variation across experiments as well as mutual nearest neighbors (Butler et al., 2018; Haghverdi et al., 2018). Integration features were selected based on the top 4000 highly variable features using “SelectIntegrationFeatures” function (nfeatures = 4000), which was used as input for the “anchor.features” argument of the “FindIntegrationAnchors” function. PCA analysis was performed on the 4000 variable features and the top 50 principal components selected based on the elbow plot heuristic, which measures the contribution of variation in each component. These 50 principal components were used in “FindNeighbors” and “FindClusters” functions to perform graph-based clustering on a shared nearest neighbor graph (Levine et al., 2015; Xu and Su, 2015). Louvain algorithm was used for modularity optimization in clustering the cells using “FindClusters” function. The resolution parameter (res = 0.4) that determines the granularity of the clustering was selected by visually inspecting clusters with resolutions ranging 0.1 - 2.0 as well as clustree graphs (Zappia and Oshlack, 2018). Uniform Manifold Approximation and Reduction (UMAP) was used for non-linear dimensional reduction of the first 50 principal components and visualize the data using “RunUMAP” function (Becht et al., 2018). Data was graphed using different plot functions, such as “DimPlot”, “VlnPlot”, “FeaturePlot”, “Dotplot” and “DoHeatmap”, to view the cell cluster identity and marker gene expression. Cell proportion data was extracted using “table” and “prop.table” functions. Differential gene expression for individual cluster was identified using Wilcoxon rank sum test in the “FindAllMarkers” function. Marker genes detected in at least 25% of the clustered cells and with a logFC threshold of 0.25 were selected. Only positive markers were reported.

### Subset analysis of neuron clusters

The neuron cell clusters identified from the complete dataset were subclustered using the “subset” function for subcluster analysis. The subset was again normalized and scaled using “SCTransform” function with glmGamPoi method (Ahlmann-Eltze and Huber, 2021). Fifty principal components were used, and the resolution parameter was set to 0.6. Further downstream analysis was done as described above for the integrated analysis. The top DE markers generated for each neuron subcluster using “FindAllMarkers” function is given in table S4.

### Cluster identification using differentially expressed markers

A “CNS markers” database of previously published markers of the different cell types that comprise the vertebrate brain and/or SC tissues was compiled (Baek *et al*., 2019; Cavone *et al*., 2021; Guillemot, 2007; Haring *et al*., 2018; Hayashi *et al*., 2018; Hernandez-Miranda *et al*., 2017; Lu *et al*., 2015; Milich *et al*., 2021; Rosenberg *et al*., 2018; Rougeot *et al*., 2019; Sathyamurthy *et al*., 2018; Tambalo *et al*., 2020; Tang *et al*., 2017; Yadav *et al*., 2023; Zeisel *et al*., 2018; Zhang *et al*., 2014). For each cell cluster, every marker gene identified as a top differentially expressed (DE) marker of that cluster was cross-referenced with our compiled “CNS markers” database (Table S1), using our “DEMarkerScoring” algorithm. For every matching marker gene, one point was given to the respective cluster under the column name with the matching cell identity. Iteration over every marker gene was performed to generate a scoring matrix with varying points for each cluster against the different cell identities compiled in the “CNS markers” database (Table S2, sheet: scoring). Then, “phyper” function in R was used to calculate binomial probabilities using hypergeometric distribution for the total score obtained by each cluster against each cell identity in the database (Table S2, sheet: Binomial probability). Then, −log10 of probability values are obtained for plotting the heatmap (Table S2, sheet: −log10P). The resulting values were scaled to 0-100 and plotted as a heatmap using GraphPad prism. Each cluster was given an identity based on the maximum −log10 P score obtained in the heatmap. The top DE markers of clusters with ambiguous scores are manually searched in the literature (Zhang *et al*., 2014; Zhang et al., 2016) (https://brainrnaseq.org/) to confirm cluster identity. “RenameIdents” function was used to assign identity to each cluster. For confirmation of the assigned cluster identity, enrichment of classical markers of respective cell types were tested using Dot plot. The “DEMarkerScoring” algorithm is available here: https://github.com/MokalledLab/Zebrafish_SCI_Atlas/blob/main/s7_marker_comparison_v3.R.

### Filtering differentially expressed markers for candidate gene selection

To identify candidate genes that are uniquely expressed in N20 iNeurons, top DE markers were cross-checked and filtered against the top DE markers of all the other neuron subclusters. Enrichment and percent expression of these unique genes were then confirmed using Dot plots. Genes that were enriched or expressed in other neuronal or non-neuronal subclusters, in addition to their expression in iNeurons, were omitted. Genes that are exclusively enriched in iNeurons were selected for generating crispants.

### Cell-cell interaction assay

The R package CellChat (v1.4.0) was used to evaluate regenerative cell-cell interactions after SCI (Jin *et al*., 2021). CellChat models the probability of cell-cell communication by integrating our gene expression data with a database (CellChatDB) of known interaction between signaling ligands, receptors, and their cofactors. CellChatDB is a manually curated from literature-supported ligand-receptor interactions in human and mouse, only. Therefore, we converted the normalized RNA data matrix of our zebrafish dataset at each time point into human orthologues using Ensemble biomaRT package (Durinck et al., 2009). The following rules were applied while converting the data matrix: 1) one-to-one orthologue mapping was performed whenever possible, 2) For genes with one to several human orthologues, the corresponding zebrafish RNA data value was copied to every mapped gene in humans, 3) RNA data values were eliminated for zebrafish genes that did not have any human orthologue, 4) RNA data values of paralogous zebrafish genes were added and the cumulative data value was assigned to the human orthologue (https://github.com/MokalledLab/Zebrafish_SCI_Atlas/blob/main/Sparse_matrix_homologe_converter_082322.R). The converted RNA data was used then to create CellChat object using “createCellChat” function, followed by the recommended preprocessing functions with default parameters for the analysis of individual datasets. Truncated mean method with 10% trimmed observation was used to compute average gene expression per cell group in for the complete dataset (Fig. 2 and S2). The default trimean method was used to compute average gene expression for the CellChat analysis of neuron subset (Fig. 5 and S5). CellChatDB.human was used as the database for inferring cell-cell communication. All categories of ligand-receptor interactions in the database were used in the analysis. Communications involving less than 10 cells were excluded. Function “netAnalysis_computeCentrality” was used to calculate network centrality scores at each time point. Functions such as “netVisual_circle”, “netAnalysis_contribution”, “netVisual_aggregate”, “netVisual_bubble” and “netAnalysis_signalingRole_heatmap” were used to generate different plots used in this paper.

### Trajectory inference analysis

Pseudotime analysis was performed using CellRank (v1.5.1) package in Python (v3.8.10) (Lange et al., 2022; Rossum and Drake, 2009). Before analysis, the Seurat object was converted to an AnnData in a two-step process using the R package SeuratDisk (v0.9020). First, the Seurat object was saved as an h5Seurat file and then, converted it to an AnnData file to use as an input for pseudotime analysis using CellRank. Basic pre-processing of the AnnData file was performed with default recommended settings using the scvelo (v0.2.4) and scanpy (v1.9.1) packages. Genes expressed in at least 10 cells were included and 30 principal components and neighbors were used for preprocessing of the data. CytoTRACEkernel was used for pseudotime computation and direct KNN graph-edges to point into the direction of increasing differentiation status (Gulati *et al*., 2020). The transition matrix was calculated by setting the threshold_scheme parameter to “soft” to reconstruct cellular trajectories using VIA algorithm (Stassen et al., 2021). This transition matrix was projected onto a UMAP plot to obtain a velocity graph.

### Gene ontology

Gene ontology analysis was performed using Metascape (Zhou et al., 2019). Input and analysis species were set as D. rerio. Express analysis was performed for gene ontology. Metascape identified all statistically enriched terms (including GO biological processes, Reactome gene set and KEGG pathway), and calculated accumulative hypergeometric p-values and enrichment factors. Significant terms were hierarchically clustered into a tree based on Kappa-statistical similarities among their gene memberships. A kappa score of 0.3 was applied to cast the tree into term clusters. The most enriched term in each cluster was chosen as the representative term.

### Regeneration Score

Genes associated with the GO term “regeneration” in the database Amigo (http://amigo.geneontology.org/amigo/landing) were identified. As a result, Regeneration Associated Genes (RAG) present in the following terms were included: axon extension involved in regeneration_GO_0048677, Axon regeneration_GO_0031103, dendrite regeneration_GO_0031104, Positive regulation of axon regeneration_GO_0048680, Positive regulation of neuron projection regeneration_GO_0070572, Regeneration_GO_0031099 (Table S5). Regeneration score for each neuron cluster was calculated based on the DE expressed markers obtained through “FindAllMarkers” function. All upregulated marker genes detected in at least 25% of the clustered cells and with a logFC threshold of 0.25 were considered top DE markers. Our previously mentioned “DEMarkerScoring” algorithm was then used to cross-reference and score each cluster based on the similarity between Top DE markers and RAG. - log10P value was used to generate the UMAP plot with the regeneration score. Raw values are detailed given in Table S4 (sheet Cluster ID).

### Histology

Sixteen µm cross cryosections of paraformaldehyde fixed SC tissues were used. Tissue sections were mainly imaged using a Zeiss LSM 800 confocal microscope (Axio Imager.Z2, Objective: Plan-Apochromat 20x/0.8 M27, Detector: Multialkali-PMT) for immunofluorescence and hybridization chain reaction (HCR) RNA *in situ* hybridization. For glial bridging and axon-regrowth assay (Fig. 8D, F), images were taken using Zeiss slide scanner (Axio Scan.Z1, Objective: Plan-Apochromat 20x/0.8 M27, Imaging device: OrcaFlash) The HCR RNA *in situ* hybridization protocol was adapted from Molecular Instruments (https://www.molecularinstruments.com/hcr-ihc-protocols) (Choi et al., 2018). Briefly, tissue sections were hydrated in PBS, dehydrated stepwise into 100% ethanol, then rehydrated stepwise into PBT (0.1% Tween-20 in PBS). Sections were pre-treated either with 0.2% TritonX-100 in PBS for 5 minutes or boiled in Citrate Buffer (10mM Citric Acid, 0.05% Tween-20, pH 6.0) for 10 minutes. For blocking, sections were incubated in Hybridization buffer (Molecular Instruments) for 1 hr at 37°C. For hybridization, sections were incubated in pre-warmed DNA probe sets diluted to 0.0015 pmol/uL in Hybridization buffer for 48 hrs at 37°C. Washes were then performed with Wash Buffer (Molecular Instruments) at 37°C followed by 5x SSCT (3M NaCl, 0.3 M Sodium Citrate, 0.1% Tween-20, pH 7.0). For signal amplification, sections were incubated in Amplification buffer (Molecular Instruments) for 1 hr at room temperature. Prior to amplification, h1 and h2 hairpins were snap-cooled in individual tubes by heating to 95°C and allowing tubes to return to ambient temperature slowly. For amplification, h1 and h2 snap-cooled hairpins were mixed together and diluted 1:50 in Amplification buffer. Amplification proceeded overnight at room temperature in the dark. Samples were then washed thoroughly in 5x SSCT, 5x SSC, and PBT at room temperature before proceeding to immunohistochemistry. HCR RNA probes used in this study (*syt11a, syt11b,* and *vamp4*) were designed using a python script. Probe sets were ordered as 50 pmol opools from IDT. Oligo sequences for each probe set are provided in Table S7.

For immunohistochemistry, tissue sections were hydrated in PBT then treated with blocking agent (5% goat serum in PBT) for 1 hr at room temperature. For nuclear antigens, sections were treated with 0.2% TritonX-100 in PBT for 5 minutes and washed thoroughly in PBT prior to the blocking step. Sections were incubated overnight at 4°C with respective primary antibodies diluted in blocking agent, then washed in PBT, and treated for 1 hour in secondary antibodies diluted in blocking agent at room temperature. For EdU staining, sections are incubated with freshly prepared EdU staining solution (100mM Tris at pH 8.5 - SIGMA T6066, 1 mM CuSO4 – SIGMA C1297, 200 µM Fluorescent azide – Thermo Fischer Scientific and 100 mM Ascorbic Acid - SIGMA A5960) for 30 minutes in the dark at room temperature (Salic and Mitchison, 2008). Following washes, sections were incubated in 1 µg/mL of Hoechst (Thermo Scientific, H3570) for 3 minutes, washed in PBT and mounted in Fluoromount-G mounting media (SouthernBiotech, 0100-01). Primary antibodies used in this study were chicken anti-GFP (AVES, 1020, 1:1000), rabbit anti-dsRed (Clontech, 632496, 1:250), mouse anti-HuC/D (Invitrogen, A21271, 1:500). Secondary antibodies (Invitrogen, 1:250) used in this study were Alexa Flour 488, Alexa Flour 594, and Alexa Flour 647 goat anti-rabbit or goat anti-mouse antibodies.

### Rostral and caudal axon tracing

Two sets of axon tracing experiments were performed at 1 wpi (Mokalled *et al*., 2016). For rostral labeling, fish were anaesthetized using MS-222 and fine scissors were used to transect the cord 4 mm rostral to the lesion site. For caudal labeling, fish were anaesthetized using MS-222 and fine scissors were used to transect the cord 4 mm caudal to the lesion site. For both sets of tracing experiments, Biocytin-soaked Gelfoam Gelatin Sponge was applied at the new injury site (Gelfoam, Pfizer, cat# 09-0315-08; Biocytin, saturated solution, Sigma, Cat# B4261). Fish were euthanized 3 hours post-treatment and Biocytin was histologically detected using Alexa Fluor 594-conjugated Streptavidin (Thermo Fisher, cat# S-11227). *syt11b* HCR in situ hybridization and HuC/D staining were performed on the same sections. The numbers of Biocytin^+^ HuC/D^+^ and Biocytin^+^ HuC/D^+^ *syt11b*^+^ neurons were calculated 150, 450, and 750 μm rostral and caudal to the lesion. The numbers of Biocytin^+^ HuC/D^+^ were normalized to HuC/D^+^ neurons to estimate the extent of labeling. The numbers of Biocytin^+^ HuC/D^+^ *syt11b*^+^ neurons were then normalized to Biocytin^+^ HuC/D^+^ neurons to estimate the extent of iNeuron integration in the labeled circuitry. We note that in the axon tracing experiments performed at 1 wpi (Figure 7 and S7), we observed ectopic Biocytin labelling of ependymal glial cells on the non-labeled side of the cord distal to the lesion site (caudal ependymal glial cells distal to the lesion were Biocytin^+^ following rostral tracing; rostral ependymal glial cells distal to the lesion were Biocytin^+^ following caudal tracing). This labeling is not usually observed by axon tracing at 4 or 6 wpi.

### CRISPR/Cas9 mutagenesis

CRISPR/Cas9 design, mutagenesis and screening was performed as previously described (Klatt Shaw and Mokalled, 2021). Briefly, crRNA guide RNA sequences were selected using CHOPCHOP (https://chopchop.cbu.uib.no/). Only sequences with no off-target sites with three or fewer mismatches elsewhere in the genome were selected. To maximize the effect of small indels, target sites were chosen that lie within essential domains (Table S5). Alt-R tracrRNA and crRNA gRNAs (IDT, Cat# 1072534) were reconstituted using manufacturer’s specifications as 100 μM stocks, then annealed at a final concentration of 50 μM. Alt-R S.p. Cas9 nuclease V3 (IDT, Cat# 1081059) was diluted in Cas9 dilution buffer (1 M HEPES at pH 7.5, 2 M KCl) to a working concentration of 25 μM. Annealed dgRNA duplexes were diluted 1:1 in duplex buffer (IDT, Cat# 11-05-01-03) to a working concentration of 25 μM. Equal volumes of dgRNA were added to Cas9 protein and incubated at 37°C for 5 minutes. For dual targeting of *syt11a* and *syt11b*, Cas9 protein was added in equal molar amounts to the total concentration of dgRNA. Tubingen wild-type embryos were injected with 1 nL of CRISPR/Cas9 solution at the one-cell stage and grown to adulthood for genotyping, SC surgeries and functional analysis. The full sequence for each guide RNAs is provided in Table S6.

### Genotyping by capillary electrophoresis

Capillary electrophoresis was used to calculate the indel frequency for each CRISPR/Cas9 target site. For DNA extraction, whole 4 dpf larvae or ∼3 mm of excised adult tail fins were added to 20 μL (larvae) or 50 μL (adult fin) of fin lysis buffer (10 mM Tris at pH 7.0, 50 mM KCl, 1 mM EDTA at pH 8.0, 0.3% Tween-20, freshly added Proteinase K at 200 µg/ml). DNA samples were incubated at 55°C for 60 minutes and then, followed by a 15-minute incubation at 95°C. 150-250 bp PCR products were generated using NEB Taq Polymerase (Cat# M0273) with gene-specific primers (Table S5) in a volume of 10 μL in Pryme PCR semi-skirted PCR plates (MidSci, Cat# AVRT1). Samples were diluted to 24 μL in TE dilution buffer (Agilent, DNF-495-0060) and loaded into the 5200 Fragment Analyzer System (Agilent, Cat# M5310AA). Capillary electrophoresis was carried out using the Agilent Fragment Analyzer Qualitative DNA Kit (Cat# DNF-905-K1000) according to the manufacturer’s specifications.

To calculate indel frequency, 2 wild-type control siblings were PCR amplified. The size of wild-type products was determined using these control samples. PCR products with peaks within 1 bp of control amplicon size were termed wild types (non-indel). Indel frequency was calculated by dividing the signal intensity of total non-indel peaks by the cumulative intensity of all peaks in each sample. Because of significant noise due to primers (<100 bp) and non-specific products (>300 bp) in wild-type samples, only signals between 100 and 300 bp were used to calculate indel frequency.

### Swim endurance assay

Swim endurance assays were performed as previously described (Burris *et al*., 2021). Zebrafish were exercised in groups of 8-15 in a 5 L swim tunnel device (Loligo, Cat# SW100605L, 120V/60Hz). After 5 minutes of acclimation (with no flow) inside the enclosed tunnel, fish were swum at 9 cm/s and 10 cm/s for 5 minutes each to evaluate their swimming capability at bare minimum velocity. Water current velocity was then increased every two minutes and fish swam against the current until they reached exhaustion. Exhausted animals were removed from the chamber without disturbing the remaining fish. Swim time at exhaustion was recorded for each fish. Fish that were exhausted before the first 15 minutes were considered poorly recovered. Percent recovery was calculated by dividing the number of fish that swam more than 15 minutes to the total number of fish present in the group.

### Swim behavior assay

Swim behavior assays were performed as previously described (Burris *et al*., 2021). Zebrafish were divided into groups of 5 in a 5 L swim tunnel device (Loligo, Cat# SW100605L, 120V/60Hz). Each group was allowed to swim for a total of 15 min under zero to low current velocities (5 min at 0 cm/s, 5 min at 10 cm/s, and 5 min at 15 cm/s). The entire swim session was recorded using high-speed camera (iDS, USB 3.0 color video camera) with following settings: aspect ratio, 1:4; pixel clock, 344; frame rate, 70 frames/s; exposure time: 0.29; aperture, 1.4 to 2; maximum frames; 63,000. Movies were converted to 20 frames/s and analyzed using a customized Fiji macro (https://github.com/MokalledLab/Zebrafish_SCI_Atlas/blob/main/Swim%20behavior_Tracking_v2.ijm). For each frame, animals/objects >1500 px2 were identified, and the XY coordinates were derived for each animal/object. Each frame was independently analyzed, and animal/object tracking was completed using a customized R Studio script (https://github.com/MokalledLab/Zebrafish_SCI_Atlas/blob/main/SwimBehavior_processing%20and%20quantification_v7.R). The script aligned coordinates, calculated swim metrics considering three separate frame windows (Frames 0-6000 at 0 cm/s; frames 6001-12000 at 10 cm/s, and frames 12001-18001 at 20 cm/s). Distance swam was analyzed at 0, 10, and at 15 cm/s.

### Axon regrowth assay

Anterograde axon tracing was performed on adult zebrafish at 28 dpi as previously described (Mokalled *et al*., 2016). Fish were anaesthetized using MS-222 and fine scissors were used to transect the cord 4 mm rostral to the lesion site. Biocytin-soaked Gelfoam Gelatin Sponge was applied at the new injury site (Gelfoam, Pfizer, cat# 09-0315-08; Biocytin, saturated solution, Sigma, Cat# B4261). Fish were euthanized 3 hours post-treatment and Biocytin was histologically detected using Alexa Fluor 594-conjugated Streptavidin (Thermo Fisher, cat# S-11227). Biocytin-labeled axons were quantified using the “threshold” and “particle analysis” tools in the Fiji software. Four sections per fish at 0.5 (proximal) and 2 (distal) mm caudal to the lesion core, and 2 sections 1 mm rostral to the lesion, were analyzed. Axon growth was normalized to the efficiency of Biocytin labeling rostral to the lesion for each fish. The axon growth index was then normalized to the control group for each experiment.

### Glial bridging assay

Gfap immunohistochemistry was performed on serial sections. The cross-sectional area of the glial bridge and the area of the intact SC rostral to the lesion were measured using ImageJ software. Bridging was calculated as a ratio of these measurements.

### Quantifications

#### Cell counting

Cell counting were performed using a customized Fiji script (adapting ITCN plugin: Image based Tool for counting nuclei - https://github.com/MokalledLab/Zebrafish_SCI_Atlas/blob/main/Cell_Counter_v1-5_vms.ijm).

Orthogonal projections of individual image stacks were generated using Zen software. A customized Fiji script incorporated user-defined inputs to define channels (including Hoechst) and to outline SC perimeters. To quantify nuclei, the following parameters were set in ITCN counter: width – 15; minimal distance – 7.5; threshold – 0.4. Once nuclei were identified, user-defined thresholds of individual cell markers were used to mask the image and identify nuclei located inside the masked regions. X/Y coordinates were extracted for each nucleus for cell counting. Raw counts and X/Y coordinates from Fiji were processed using a customized R script (https://github.com/MokalledLab/Zebrafish_SCI_Atlas/blob/main/Cell_Count_Processor_v1_vms.R). Markers that share nuclei with the same X/Y coordinates were considered overlapping.

#### Quantifying HCR in situ signal

HCR *in situ* hybridization signals were quantified using a customized Fiji script (https://github.com/MokalledLab/Zebrafish_SCI_Atlas/blob/main/hcr_celltypespecific_analysis_v 3.1_080222.ijm). Orthogonal projections of individual image stacks were generated using the Zen software (Zeiss). ROIs around SC borders were defined to exclude non-SC tissues and user-defined thresholds were set to create an inverted mask. Background noise was either manually corrected or defined by thresholding a dedicated, unstained background channel. Integrated density was quantified for fluorescence signals inside the quantification mask using the “Analyze Particles” command. To calculate the HCR *in situ* hybridization signal inside neurons, an inverted mask was created after thresholding HuC/D^+^ neurons, then added to the final quantification mask. The fluorescence signals inside this newly generated mask were considered neuron specific.

