## Supplemental Figures for "Single-cell analysis of innate spinal cord regeneration identifies intersecting modes of neuronal repair"

**Figure S1**

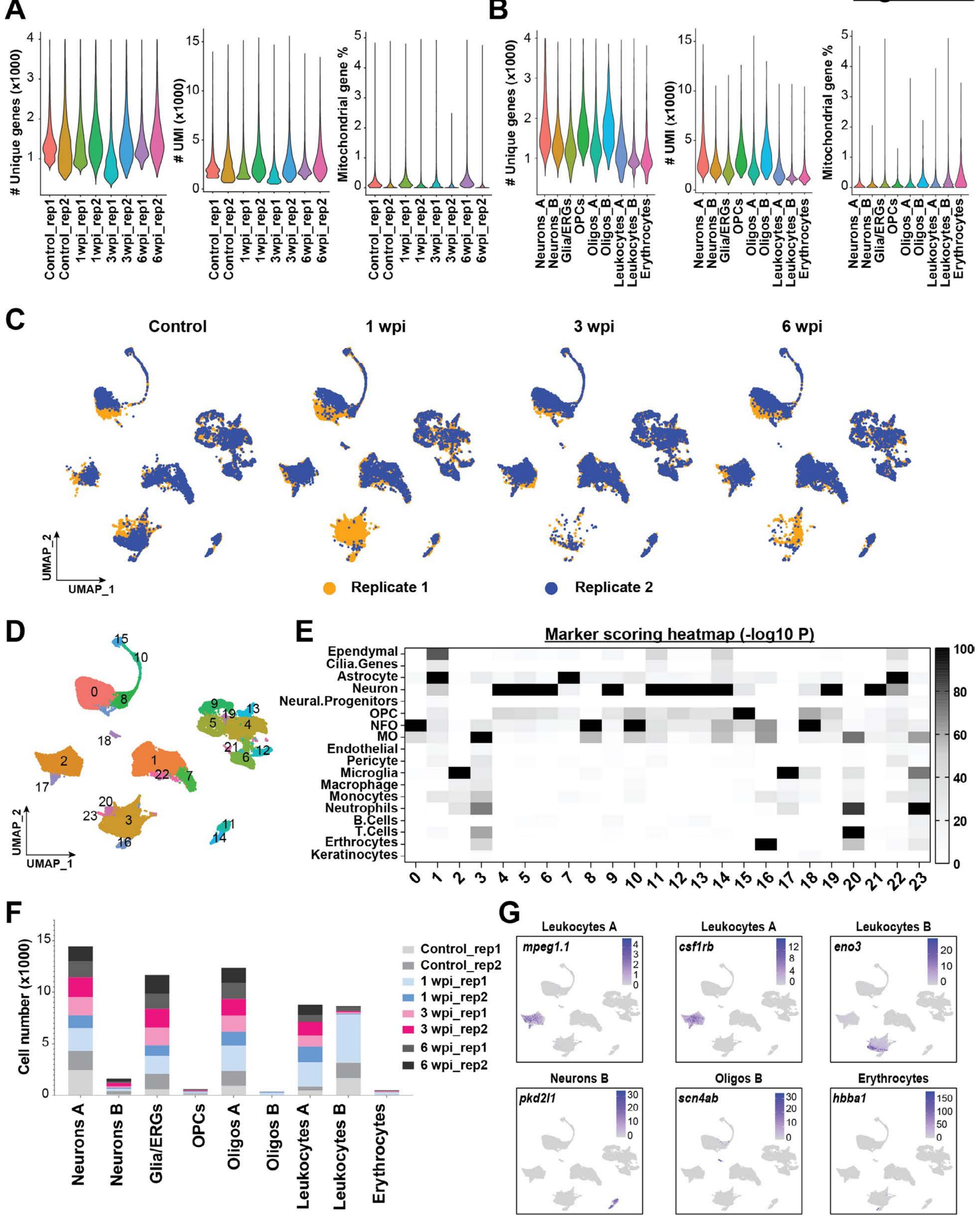
A
### Unique genes (x1000)
### UMI (x1000)
Mitochondrial gene %
Control_rep1
Control_rep2
1wpi_rep1
1wpi_rep2
3wpi_rep1
3wpi_rep2
6wpi_rep1
6wpi_rep2
B
Neurons_A
Neurons_B
Glia/ERGs
OPCs
Oligos_A
Oligos_B
Leukocytes_A
Leukocytes_B
Erythrocytes
C
Control
1 wpi
3 wpi
6 wpi
UMAP_2
UMAP_1
Replicate 1
Replicate 2
D
E
Marker scoring heatmap (-log10 P)
Ependymal
Cilia.Genes
Astrocyte
Neuron
Neural.Progenitors
OPC
NFO
MO
Endothelial
Pericyte
Microglia
Macrophage
Monocytes
Neutrophils
B.Cells
T.Cells
Erthrocytes
Keratinocytes
F
Cell number (x1000)
Control_rep1
Control_rep2
1 wpi_rep1
1 wpi_rep2
3 wpi_rep1
3 wpi_rep2
6 wpi_rep1
6 wpi_rep2
Neurons A
Neurons B
Glia/ERGs
OPCs
Oligos A
Oligos B
Leukocytes A
Leukocytes B
Erythrocytes
G
Leukocytes A
mpeg1.1
Leukocytes A
csf1rb
Leukocytes B
eno3
Neurons B
pkd2l1
Oligos B
scn4ab
Erythrocytes
hbba1

**Figure S1. Quality control metrics, cluster identification of snRNA-seq data and raw cell counts (Related to figure 1). (A,B)** Violin plots showing the distribution of cells based on their quality control metrics grouped by both replicates and cell types. Only cells that are retained after removal of low-quality cells are shown. The total number of unique genes found per cell, the number of unique molecular identifiers (UMI) per cell, and the proportion of UMIs that map to mitochondrial protein encoding genes per cell are shown. **(C)** UMAP plot represents the overlap between the two replicate pools at each time point, with 50 SC tissues pooled within each replicate. The number of cells present in each replicate at different time point are control (6432, 6965), 1 wpi (14312, 5416), 3 wpi (6533, 7211) and 6 wpi (5848, 6256). **(D)** Merged UMAP plot of the complete dataset showing 24 coarse clusters identified at 0.4 resolution parameter. **(E)** Heatmap depicting the -log10 P value for the marker scoring output from "DEMarkerScoring" algorithm after cross-referencing the top DE markers of each cluster with our assembled VNM marker database. Hypergeometric probability test was used to calculate the P value. P values are detailed in Table S2. -log10 P value for each cluster is scaled between 0-100. Cluster identity is predicted based on the maximum -log10 P value obtained, which represents the darkest quadrant in the heatmap. **(F)** Bar graph showing the number of cells that mapped to different cell populations split by biological replicate across the time course of regeneration. **(G)** Feature plots showing the distributions of canonical marker genes corresponding to major cell populations.

### Figure S2

**A**

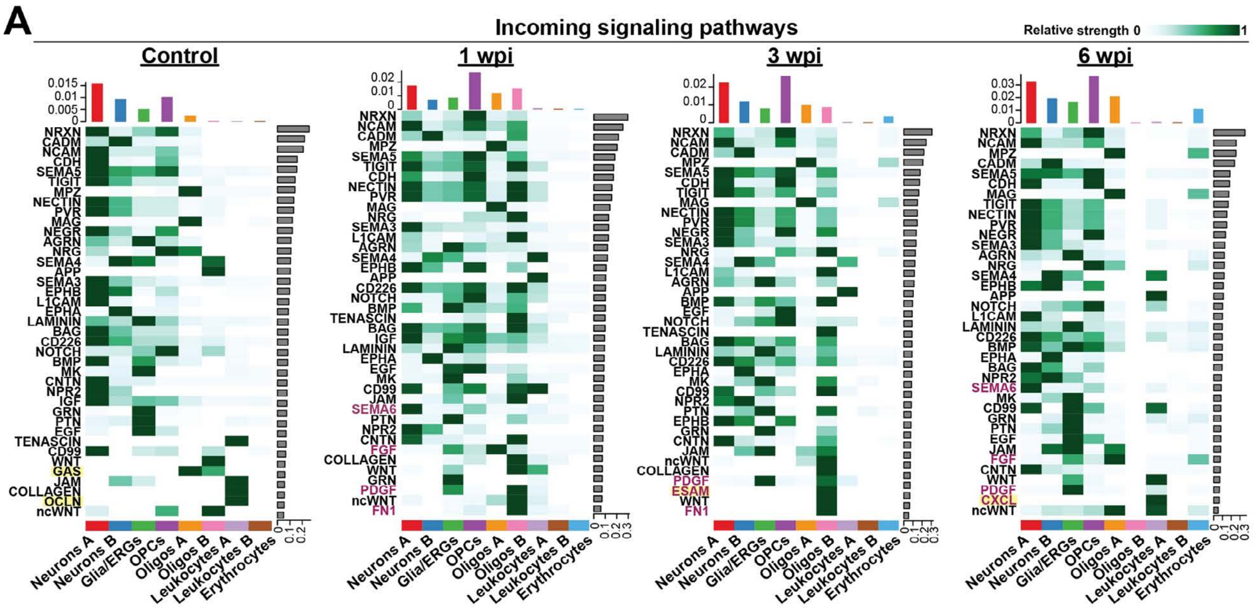


**B**

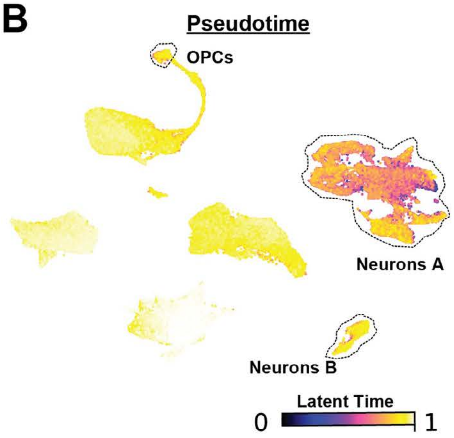


**C**

**Incoming signals to neurons**

| Rank | Control | 1 wpi | 3 wpi | 6 wpi |
|---|---|---|---|---|
| 1 | NRXN | NRXN | NRXN | NRXN |
| 2 | CADM | NCAM | NCAM | NCAM |
| 3 | NCAM | CADM | CADM | MPZ |
| 4 | CDH | MPZ | MPZ | CADM |
| 5 | SEMA5 | SEMA5 | SEMA5 | SEMA5 |
| 6 | TIGIT | TIGIT | CDH | CDH |
| 7 | MPZ | CDH | TIGIT | MAG |
| 8 | NECTIN | NECTIN | MAG | TIGIT |
| 9 | PVR | PVR | NECTIN | NECTIN |
| 10 | MAG | MAG | PVR | PVR |
| 11 | NEGR | NRG | NEGR | NEGR |
| 12 | AGRN | SEMA3 | SEMA3 | SEMA3 |
| 13 | NRG | L1CAM | NRG | AGRN |
| 14 | SEMA4 | AGRN | SEMA4 | NRG |
| 15 | SEMA3 | SEMA4 | L1CAM | SEMA4 |
| 16 | EPHB | EPHB | AGRN | EPHB |
| 17 | L1CAM | CD226 | BMP | NOTCH |
| 18 | EPHA | NOTCH | EGF | L1CAM |
| 19 | LAMININ | BMP | NOTCH | LAMININ |
| 20 | BAG | BAG | BAG | CD226 |
| 21 | CD226 | IGF | LAMININ | BMP |
| 22 | NOTCH | LAMININ | CD226 | EPHA |
| 23 | BMP | EPHA | EPHA | BAG |
| 24 | MK | EGF | MK | NPR2 |
| 25 | CNTN | MK | CD99 | SEMA6 |
| 26 | NPR2 | CD99 | NPR2 | MK |
| 27 | IGF | JAM | PTN | CD99 |
| 28 | GRN | SEMA6 | EPHB | GRN |
| 29 | PTN | PTN | CNTN | PTN |
| 30 | EGF | NPR2 | JAM | EGF |
| 31 | CD99 | CNTN | ncWNT | JAM |
| 32 | JAM | FGF | PDGF | FGF |
| 33 | ncWNT | WNT | | CNTN |
| 34 | | ncWNT | | PDGF |
| 35 | | | | ncWNT |

**Outgoing signals from neurons**

| Rank | Control | 1 wpi | 3 wpi | 6 wpi |
|---|---|---|---|---|
| 1 | NRXN | NRXN | NRXN | NRXN |
| 2 | CADM | NCAM | NCAM | NCAM |
| 3 | NCAM | CADM | CADM | MPZ |
| 4 | CDH | MPZ | MPZ | CADM |
| 5 | SEMA5 | SEMA5 | SEMA5 | SEMA5 |
| 6 | TIGIT | TIGIT | CDH | CDH |
| 7 | MPZ | CDH | TIGIT | MAG |
| 8 | NECTIN | NECTIN | MAG | TIGIT |
| 9 | PVR | PVR | NECTIN | NECTIN |
| 10 | MAG | MAG | PVR | PVR |
| 11 | NEGR | NRG | NEGR | NEGR |
| 12 | AGRN | SEMA3 | SEMA3 | SEMA3 |
| 13 | NRG | L1CAM | NRG | AGRN |
| 14 | SEMA4 | AGRN | SEMA4 | NRG |
| 15 | APP | SEMA4 | L1CAM | SEMA4 |
| 16 | SEMA3 | EPHB | AGRN | EPHB |
| 17 | EPHB | APP | APP | APP |
| 18 | L1CAM | NOTCH | EGF | NOTCH |
| 19 | EPHA | TENASCIN | NOTCH | L1CAM |
| 20 | LAMININ | BAG | TENASCIN | LAMININ |
| 21 | BAG | LAMININ | BAG | EPHA |
| 22 | NOTCH | EPHA | LAMININ | BAG |
| 23 | MK | EGF | EPHA | NPR2 |
| 24 | CNTN | MK | MK | MK |
| 25 | NPR2 | JAM | NPR2 | PTN |
| 26 | PTN | PTN | PTN | EGF |
| 27 | EGF | NPR2 | EPHB | JAM |
| 28 | TENASCIN | CNTN | CNTN | FGF |
| 29 | CD99 | FGF | JAM | CNTN |
| 30 | WNT | FN1 | | |
| 31 | GAS | | | |
| 32 | JAM | | | |
| 33 | | | | |
| 34 | | | | |
| 35 | | | | |

Uniquely enriched at the timepoint

Injury-induced signaling pathways

**Figure S2. Injury-induced signaling pathways and pseudotime analysis (Related to figure 2). (A)** Heatmap representation of the relative strengths of incoming signaling pathways in coarse cell populations at 0, 1, 3 and 6 wpi. Bar graphs at the top of each heatmap show cumulative signaling strengths per cell population. Bar graphs at the right of each heatmap show cumulative signaling strength per pathway. Pathways highlighted in yellow are specifically enriched at one time point. Pathways highlighted in magenta are enriched after injury. **(B)** UMAP plot with embedded pseudotime score was generated using the CytoTRACE algorithm in CellRank. Pseudotime scores are color-coded, with yellow dots representing cells with more differentiated status and blue dots representing cells with progenitor-like transcriptional status. **(C)** Tables show lists of incoming and outgoing signaling pathways activated and received by neurons based on CellChat analysis of the integrated dataset. Pathways are ranked in the descending order of their cumulative signal strengths. Pathways highlighted in magenta represent injury-enriched signaling. Pathways highlighted in magenta and yellow represent signaling pathways that are specifically enriched at the respective timepoint.

**Figure S3**

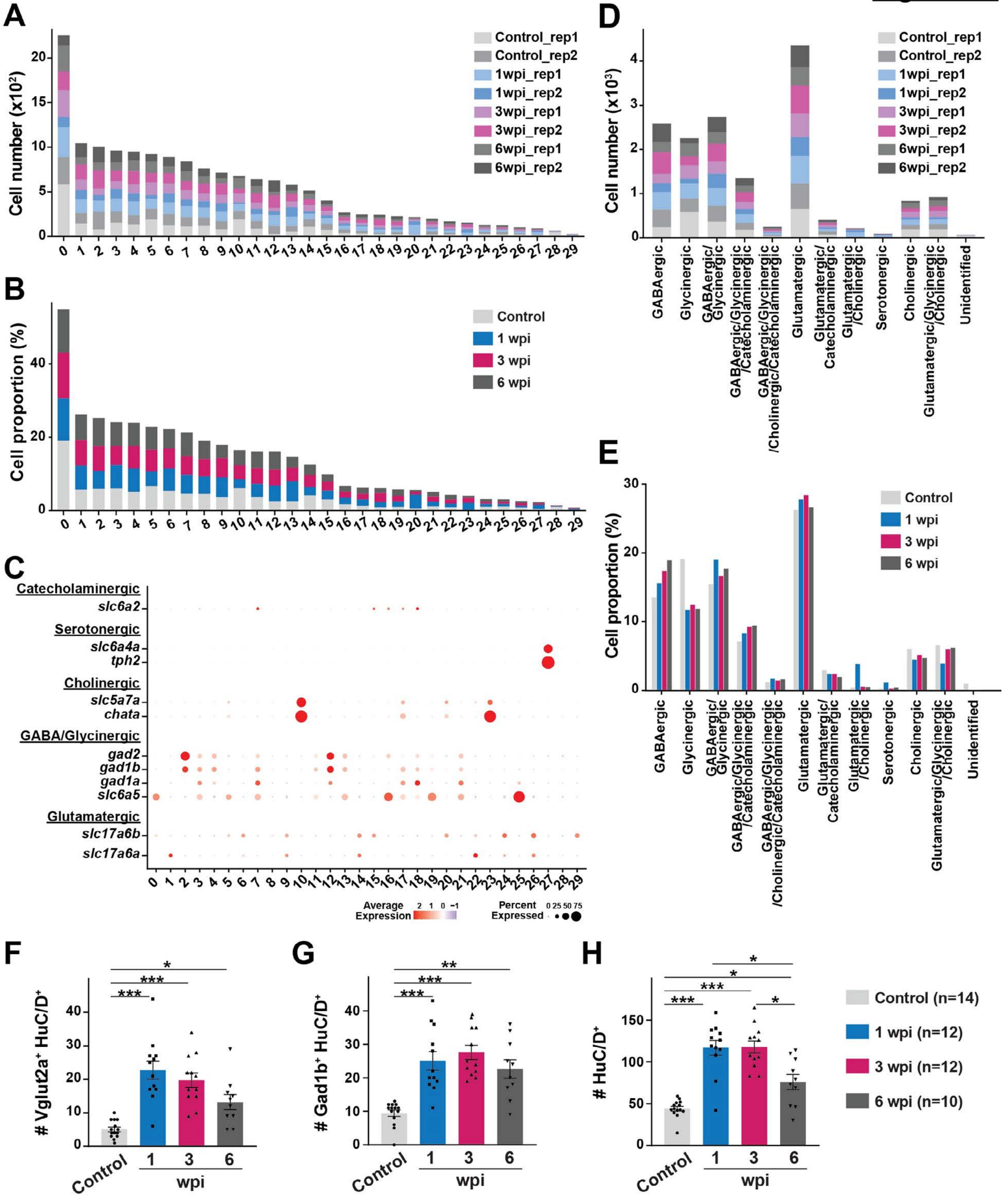
A
Cell number (x10²)
Control_rep1
Control_rep2
1wpi_rep1
1wpi_rep2
3wpi_rep1
3wpi_rep2
6wpi_rep1
6wpi_rep2
B
Cell proportion (%)
Control
1 wpi
3 wpi
6 wpi
C
Catecholaminergic
slc6a2
Serotonergic
slc6a4a
tph2
Cholinergic
slc5a7a
chata
GABA/Glycinergic
gad2
gad1b
gad1a
slc6a5
Glutamatergic
slc17a6b
slc17a6a
Average Expression
Percent Expressed
D
Cell number (x10³)
GABAergic
Glycinergic
GABAergic/Glycinergic
GABAergic/Glycinergic/Catecholaminergic
GABAergic/Glycinergic/Cholinergic/Catecholaminergic
Glutamatergic
Glutamatergic/Catecholaminergic
Glutamatergic/Cholinergic
Serotonergic
Cholinergic
Glutamatergic/Glycinergic/Cholinergic
Unidentified
E
Cell proportion (%)
F
### Vglut2a⁺ HuC/D⁺
Control
wpi
G
### Gad1b⁺ HuC/D⁺
H
### HuC/D⁺
Control (n=14)
1 wpi (n=12)
3 wpi (n=12)
6 wpi (n=10)

**Figure S3. Neurotransmitter gene expression, cell proportion and top DE markers in neuron subclusters after SCI (Related to figure 3). (A-B)** Split bar charts show the numbers and proportions of neurons that mapped to each neuron subcluster. Cell proportions for each cluster and time point in B were normalized to the total number of neurons analyzed at that time point. **(C)** Expression of canonical neurotransmitter genes was used to classify excitatory, inhibitory, cholinergic, serotonergic and catecholaminergic neuron populations. Clusters that show enrichment (gradient of red dots) for any of these marker genes are given a respective neurotransmitter identity. **(D-E)** The total numbers and proportions of different neuron subpopulations based on their neurotransmitter properties at 0, 1, 3 and 6 wpi. Cell proportions for each cluster and time point in E were normalized to the total number of neurons analyzed at that time point. **(F-H)** Quantification of glutamatergic and GABAergic neurons after SCI. Absolute numbers of *vglut2a*$^+$ HuC/D$^+$, *gad1b*$^+$ HuC/D$^+$ and HuC/D$^+$ neurons are shown. SC cross-sections at 0, 1, 3 and 6 wpi were used for quantification. Brown-Forsythe and Welch ANOVA test was performed on F, G and H. ANOVA p-value is <0.0001 for F-H. Dunnett's T3 multiple comparisons test was performed across different timepoints with 95% CI. * p≤0.05, ** p ≤0.01, *** p<0.001. SC cross sections 450 μm rostral to the lesion were analyzed in F-H. Solid circles indicate individual animals and sample sizes are indicated in parentheses.

#### Figure S4

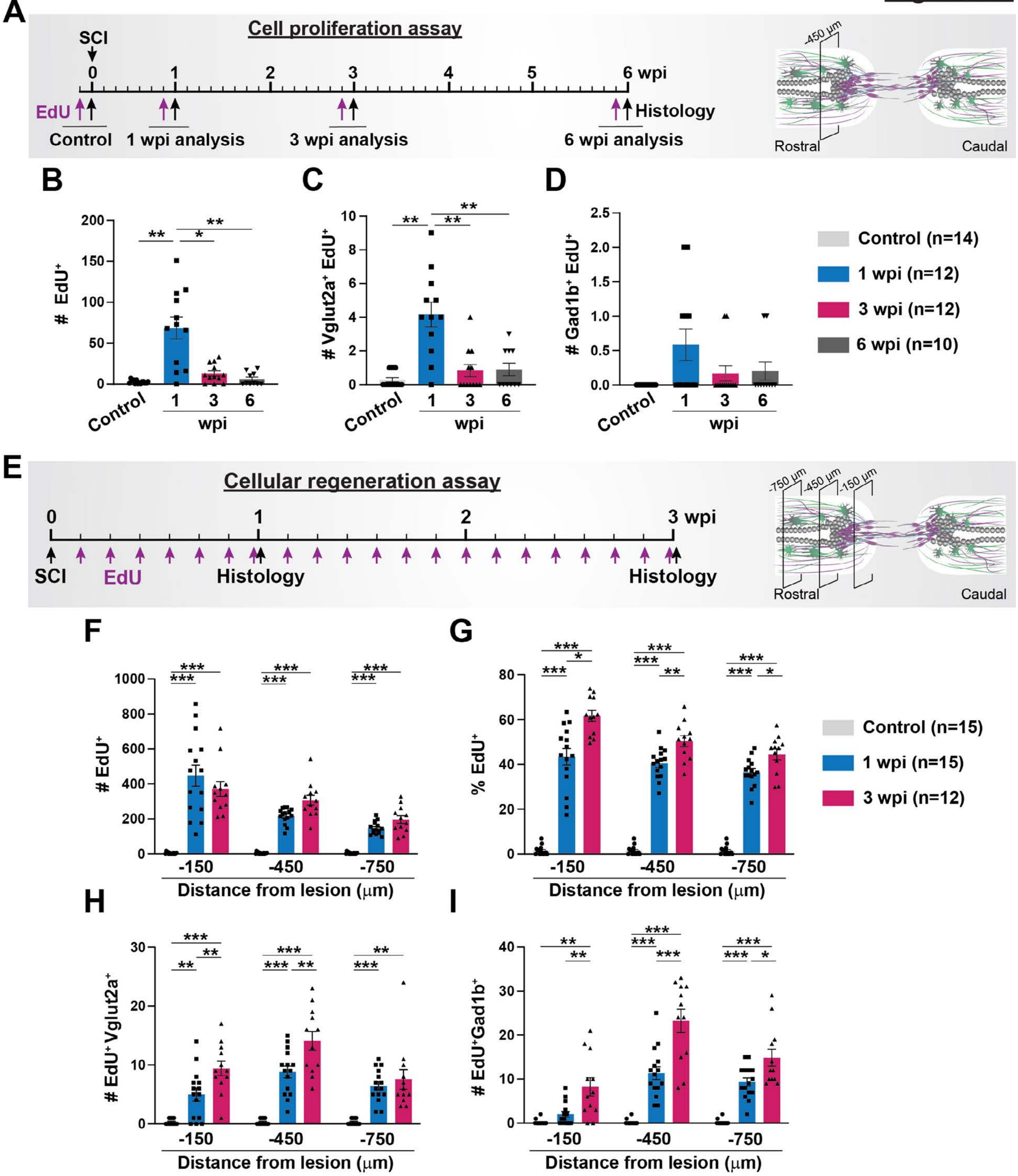

A
Cell proliferation assay
SCI
0 1 2 3 4 5 6 wpi
EdU
Histology
Control
1 wpi analysis
3 wpi analysis
6 wpi analysis
-450 µm
Rostral
Caudal
B
### EdU+
Control
1 3 6
wpi
C
### Vglut2a+ EdU+
D
### Gad1b+ EdU+
Control (n=14)
1 wpi (n=12)
3 wpi (n=12)
6 wpi (n=10)
E
Cellular regeneration assay
0 1 2 3 wpi
SCI
EdU
Histology
-750 µm
-450 µm
-150 µm
F
### EdU+
G
% EdU+
Control (n=15)
1 wpi (n=15)
3 wpi (n=12)
H
### EdU+Vglut2a+
I
### EdU+Gad1b+
-150 -450 -750
Distance from lesion (µm)

**Figure S4. Quantification of regenerating excitatory and inhibitory neurons after SCI (Related to figure 4). (A)** Experimental timeline to assess cell proliferation at 1, 3 and 6 wpi. Tg(*vglut2a*:RFP; *gad1b*:GFP) zebrafish were subjected to SC transections. Single intraperitoneal EdU injections were performed at either 6, 20, or 41 days post-injury. SC tissues were harvested for analysis at 1, 3, or 6 wpi and at 24 hours after EdU injection. Uninjured animals were injected with EdU and collected 24 hours after EdU injection as controls. SC cross-section 450 μm rostral to the lesion was used for quantifications. **(B-D)** Shown are the absolute numbers of $EdU^+$ cells (ANOVA p-value <0.0001), newly formed glutamatergic neurons ($vglut2a^+$ $EdU^+$) (ANOVA p-value <0.0001), and GABAergic neurons ($gad1b^+$ $EdU^+$) (ANOVA p-value: 0.0445). **(E)** Experimental timeline to assess the cumulative profiles of regenerating neurons at 1 and 3 wpi. Tg(*vglut2a*:RFP; *gad1b*:GFP) zebrafish were subjected to SC transections and daily intraperitoneal EdU injections. SC tissues were harvested for analysis at 1 and 3 wpi. Control SCs received daily EdU injection for 7 days before collection. Tissue sections 150, 450 and 750 μm rostral to the lesion were analyzed. **(F-G)** The absolute numbers and proportions of $EdU^+$ cells in SC sections are shown at 0, 1 and 3 wpi. **(H-I)** Absolute numbers newly formed glutamatergic ($vglut2a^+$ $EdU^+$) and GABAergic ($gad1b^+$ $EdU^+$) neurons in SC cross sections at 0, 1 and 3 wpi. The p-value across time point is <0.0001 for F, G, H, and I. Brown-Forsythe and Welch ANOVA test was performed on B, C and D with Dunnett's T3 multiple comparisons test performed across different time points having 95% CI. Two-way ANOVA was performed on F, G, H, and I with Tukey's multiple comparison test having 95% CI. SC cross sections 150, 450 and 750 μm rostral to the lesion were analyzed. Solid circles and polygons indicate individual animals and sample sizes are indicated in parentheses. * p≤0.05, ** p ≤0.01, *** p≤0.001.

### Figure S5

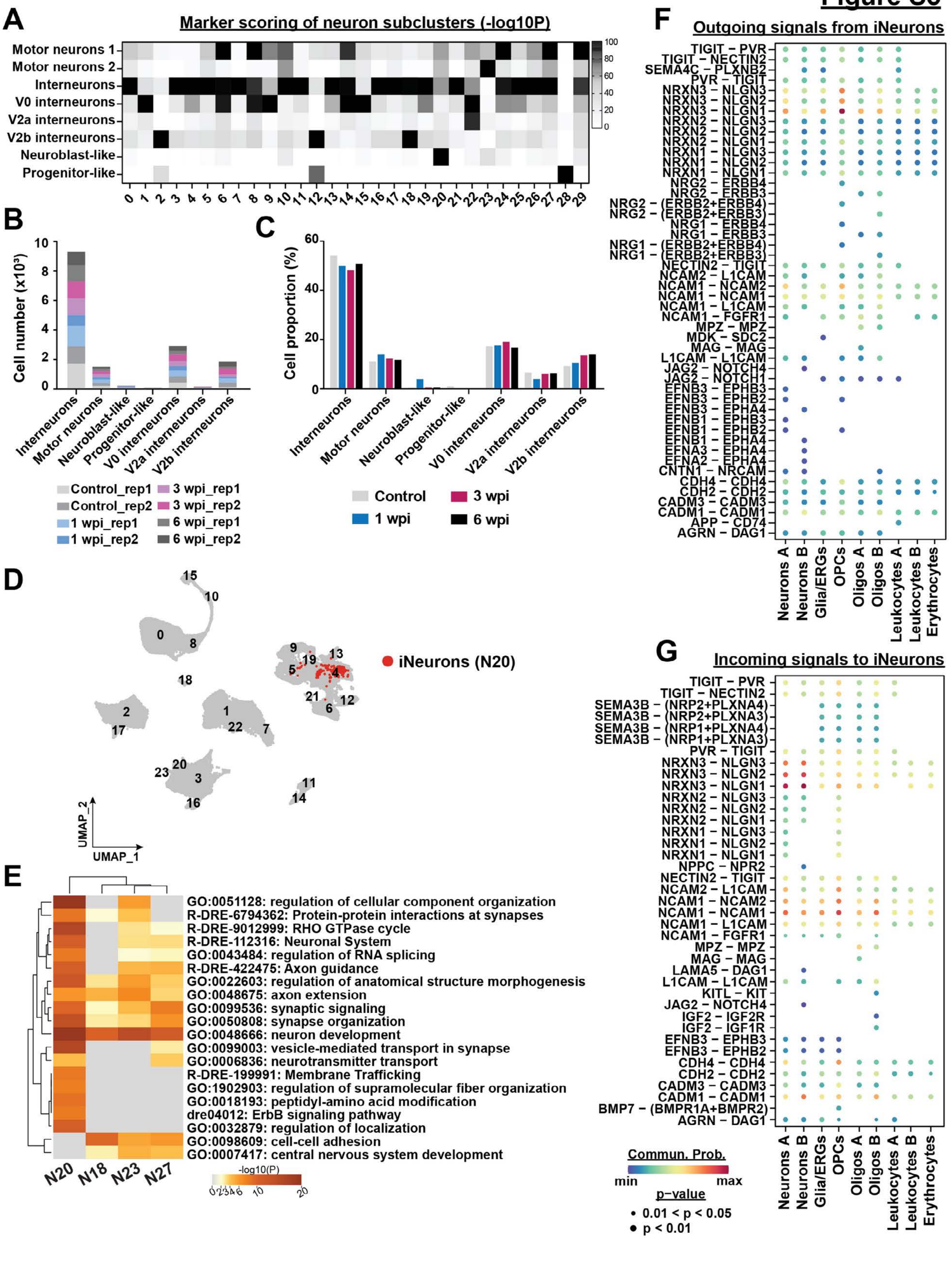

A
Marker scoring of neuron subclusters (-log10P)
Motor neurons 1
Motor neurons 2
Interneurons
V0 interneurons
V2a interneurons
V2b interneurons
Neuroblast-like
Progenitor-like
0 1 2 3 4 5 6 7 8 9 10 11 12 13 14 15 16 17 18 19 20 21 22 23 24 25 26 27 28 29
100 80 60 40 20 0
B
Cell number (x10³)
Interneurons
Motor neurons
Neuroblast-like
Progenitor-like
V0 interneurons
V2a interneurons
V2b interneurons
Control_rep1
Control_rep2
1 wpi_rep1
1 wpi_rep2
3 wpi_rep1
3 wpi_rep2
6 wpi_rep1
6 wpi_rep2
C
Cell proportion (%)
Control
1 wpi
3 wpi
6 wpi
D
iNeurons (N20)
UMAP_1
UMAP_2
E
GO:0051128: regulation of cellular component organization
R-DRE-6794362: Protein-protein interactions at synapses
R-DRE-9012999: RHO GTPase cycle
R-DRE-112316: Neuronal System
GO:0043484: regulation of RNA splicing
R-DRE-422475: Axon guidance
GO:0022603: regulation of anatomical structure morphogenesis
GO:0048675: axon extension
GO:0099536: synaptic signaling
GO:0050808: synapse organization
GO:0048666: neuron development
GO:0099003: vesicle-mediated transport in synapse
GO:0006836: neurotransmitter transport
R-DRE-199991: Membrane Trafficking
GO:1902903: regulation of supramolecular fiber organization
GO:0018193: peptidyl-amino acid modification
dre04012: ErbB signaling pathway
GO:0032879: regulation of localization
GO:0098609: cell-cell adhesion
GO:0007417: central nervous system development
N20 N18 N23 N27
-log10(P)
F
Outgoing signals from iNeurons
TIGIT – PVR
TIGIT – NECTIN2
SEMA4C – PLXNB2
PVR – TIGIT
NRXN3 – NLGN3
NRXN3 – NLGN2
NRXN3 – NLGN1
NRXN2 – NLGN3
NRXN2 – NLGN2
NRXN2 – NLGN1
NRXN1 – NLGN3
NRXN1 – NLGN2
NRXN1 – NLGN1
NRG2 – ERBB4
NRG2 – ERBB3
NRG2 – (ERBB2+ERBB4)
NRG2 – (ERBB2+ERBB3)
NRG1 – ERBB4
NRG1 – ERBB3
NRG1 – (ERBB2+ERBB4)
NRG1 – (ERBB2+ERBB3)
NECTIN2 – TIGIT
NCAM2 – L1CAM
NCAM1 – NCAM2
NCAM1 – NCAM1
NCAM1 – L1CAM
NCAM1 – FGFR1
MPZ – MPZ
MDK – SDC2
MAG – MAG
L1CAM – L1CAM
JAG2 – NOTCH4
JAG2 – NOTCH1
EFNB3 – EPHB3
EFNB3 – EPHB2
EFNB3 – EPHA4
EFNB1 – EPHB3
EFNB1 – EPHB2
EFNB1 – EPHA4
EFNA3 – EPHA4
EFNA2 – EPHA4
CNTN1 – NRCAM
CDH4 – CDH4
CDH2 – CDH2
CADM3 – CADM3
CADM1 – CADM1
APP – CD74
AGRN – DAG1
Neurons A
Neurons B
Glia/ERGs
OPCs
Oligos A
Oligos B
Leukocytes A
Leukocytes B
Erythrocytes
G
Incoming signals to iNeurons
TIGIT – PVR
TIGIT – NECTIN2
SEMA3B – (NRP2+PLXNA4)
SEMA3B – (NRP2+PLXNA3)
SEMA3B – (NRP1+PLXNA4)
SEMA3B – (NRP1+PLXNA3)
PVR – TIGIT
NRXN3 – NLGN3
NRXN3 – NLGN2
NRXN3 – NLGN1
NRXN2 – NLGN3
NRXN2 – NLGN2
NRXN2 – NLGN1
NRXN1 – NLGN3
NRXN1 – NLGN2
NRXN1 – NLGN1
NPPC – NPR2
NECTIN2 – TIGIT
NCAM2 – L1CAM
NCAM1 – NCAM2
NCAM1 – NCAM1
NCAM1 – L1CAM
NCAM1 – FGFR1
MPZ – MPZ
MAG – MAG
LAMA5 – DAG1
L1CAM – L1CAM
KITL – KIT
JAG2 – NOTCH4
IGF2 – IGF2R
IGF2 – IGF1R
EFNB3 – EPHB3
EFNB3 – EPHB2
CDH4 – CDH4
CDH2 – CDH2
CADM3 – CADM3
CADM1 – CADM1
BMP7 – (BMPR1A+BMPR2)
AGRN – DAG1
Neurons A
Neurons B
Glia/ERGs
OPCs
Oligos A
Oligos B
Leukocytes A
Leukocytes B
Erythrocytes
Commun. Prob.
min
max
p-value
0.01 < p < 0.05
p < 0.01

**Figure S5. Cluster identification and proportion of neuron subclusters (Related to figure 5). (A)** Heatmap depicting -log10 P value for the marker scoring output obtained from the “DEMarkerScoring” algorithm. Scoring was based on cross-referencing the top DE markers for each neuron cluster with the DE markers identified in *Cavone et al, 2021*. Hypergeometric probability test was used to calculate the P value. P values are detailed in the Table S8. -log10 P value for each cluster is scaled between 0-100. Cluster identity is predicted based on the maximum -log10 P value obtained, which represents the darkest quadrant in the heatmap. **(B-C)** The total numbers and proportions of different neuron subtypes at 0, 1, 3 and 6 wpi. Cell proportions for each cluster and time point in C were normalized to the total number of neurons analyzed at that time point. **(D)** UMAP plot showing iNeurons (cluster N20 cells) within the complete dataset. **(E)** Metascape gene ontology analysis for neuron clusters N18, 20, 23 and 27. Twenty of the most enriched terms are shown. The color gradient represents -log10(p-value) for each term and cluster. **(F-G)** Bubble plots showing significant Ligand-Receptor interactions for outgoing and incoming signaling pathways that are activated or received by iNeurons at 1 wpi. Dot colors and diameters represent interaction probabilities and p-values, respectively.

Figure S6

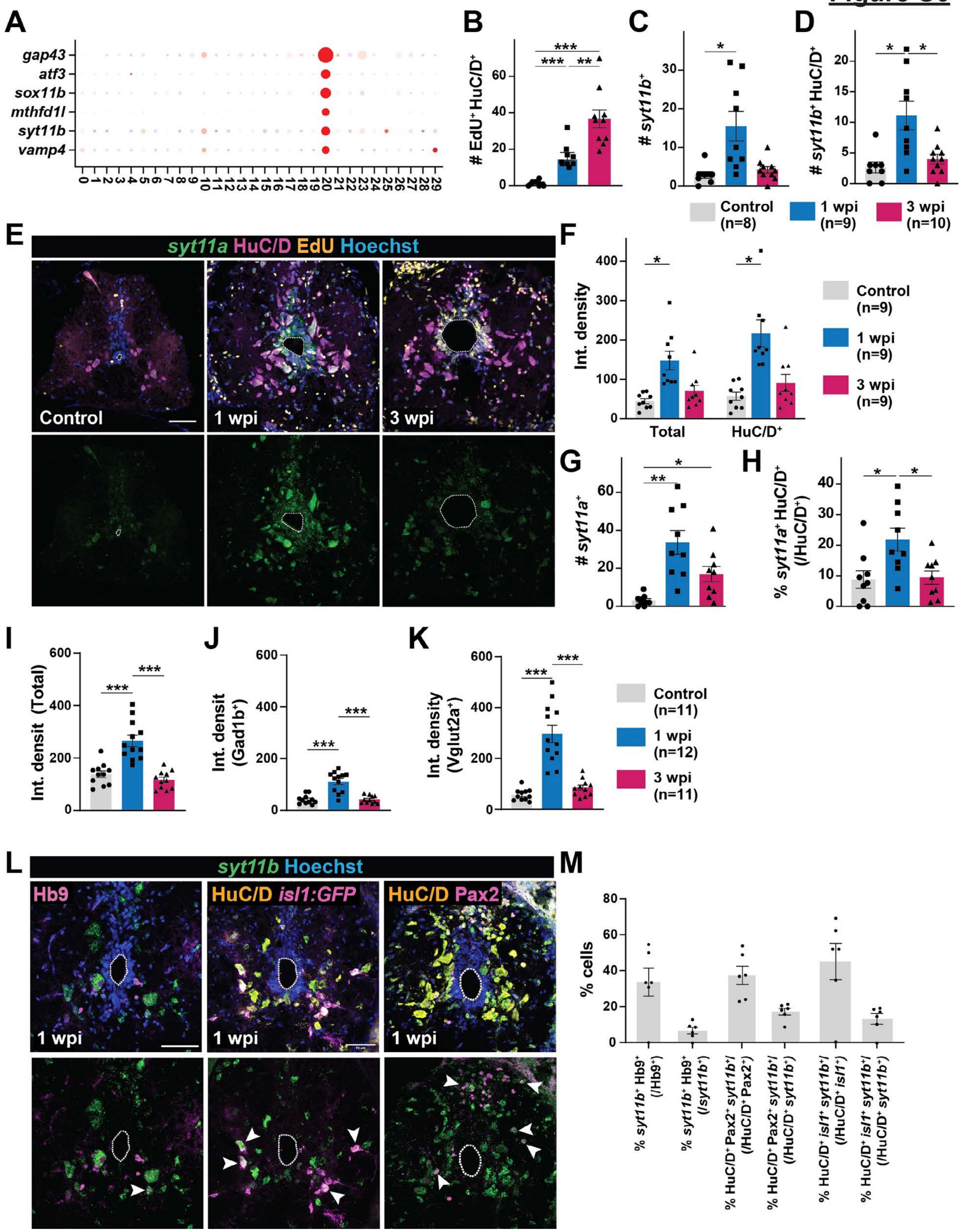
A
gap43
atf3
sox11b
mthfd1l
syt11b
vamp4
0 1 2 3 4 5 6 7 8 9 10 11 12 13 14 15 16 17 18 19 20 21 22 23 24 25 26 27 28 29
B
### EdU⁺ HuC/D⁺
C
### syt11b⁺
D
### syt11b⁺ HuC/D⁺
Control (n=8)
1 wpi (n=9)
3 wpi (n=10)
E
syt11a HuC/D EdU Hoechst
Control
1 wpi
3 wpi
F
Int. density
Total
HuC/D⁺
Control (n=9)
1 wpi (n=9)
3 wpi (n=9)
G
### syt11a⁺
H
% syt11a⁺ HuC/D⁺ (/HuC/D⁺)
I
Int. densit (Total)
J
Int. densit (Gad1b⁺)
K
Int. density (Vglut2a⁺)
Control (n=11)
1 wpi (n=12)
3 wpi (n=11)
L
syt11b Hoechst
Hb9
HuC/D isl1:GFP
HuC/D Pax2
1 wpi
M
% cells
% syt11b⁺ Hb9⁺ (/Hb9⁺)
% syt11b⁺ Hb9⁺ (/syt11b⁺)
% HuC/D⁺ Pax2⁺ syt11b⁺/ (/HuC/D⁺ Pax2⁺)
% HuC/D⁺ Pax2⁺ syt11b⁺/ (/HuC/D⁺ syt11b⁺)
% HuC/D⁺ isl1⁺ syt11b⁺/ (/HuC/D⁺ isl1⁺)
% HuC/D⁺ isl1⁺ syt11b⁺/ (/HuC/D⁺ syt11b⁺)

**Figure S6. iNeuron marker gene expression (Related to figure 6). (A)** Dot plot shows the top DE markers that were filtered based on their specific enrichment in iNeurons relative to all other spinal cell types. **(B-D)** Quantification of the total number of newly generated neurons (EdU$^+$HuC/D$^+$) (ANOVA p-value: <0.0001), *syt11b*$^+$ nuclei (ANOVA p-value: 0.0066) and *syt11b*$^+$ neurons (*syt11b*$^+$HuC/D$^+$) (ANOVA p-value: 0.0040) in whole SC cross sections at 0, 1 and 3 wpi. **(E)** HCR *in situ* hybridization for *syt11a* and staining for HuC/D, Hoechst and EdU were performed on wild-type SC cross sections at 0, 1 and 3 wpi. Dotted lines delineate central canal edges. **(F)** Quantification of *syt11a* HCR *in situ* hybridization. Integrated density was calculated in the whole SC cross sections (ANOVA p-value: 0.0013) and inside HuC/D$^+$ neurons (ANOVA p-value: 0.0008) at 0, 1 and 3 wpi. **(G-H)** Quantification of *syt11a* expressing cells after SCI. Shown are the absolute numbers of *syt11a*$^+$ cells (ANOVA p-value: 0.0008), and percent *syt11a*$^+$ neurons (*syt11a*$^+$ HuC/D$^+$ normalized to the total number of HuC/D$^+$ neurons) (ANOVA p-value: 0.0084). Analysis was performed on whole SC cross sections at 0, 1 and 3 wpi. **(I-K)** Quantification of the integrated density of *syt11b* HCR *in situ* hybridization signal in whole SC tissues (ANOVA p-value <0.0001), inside GABAergic neurons (ANOVA p-value <0.0001), and glutamatergic neurons (ANOVA p-value <0.0001). **(L)** HCR *in situ* hybridization for *syt11b* was co-localized with staining for HuC/D, Hb9, *isl1:GFP* or Pax2. Wild-type or *isl1:GFP* SC cross sections at 1 wpi were used. Dotted lines delineate central canal edges. Arrowheads point to double positive cells. **(M)** Quantification of *syt11b* expressing cells at 1 wpi. For all quantifications, solid circles and polygons in the bar charts indicate individual animals and sample sizes are indicated in parentheses. SC cross sections at 450 µm rostral to the lesion were analyzed. Brown-Forsythe and Welch ANOVA test was performed on all the graphs with Dunnett's T3 multiple comparisons test performed across time points with 95% CI. Ordinary one-way ANOVA with Tukey's multiple comparisons test (95% CI) was performed in H. * p≤0.05, ** p≤0.01, *** p≤0.001. Scale bars, 50 µm.

### Figure S7

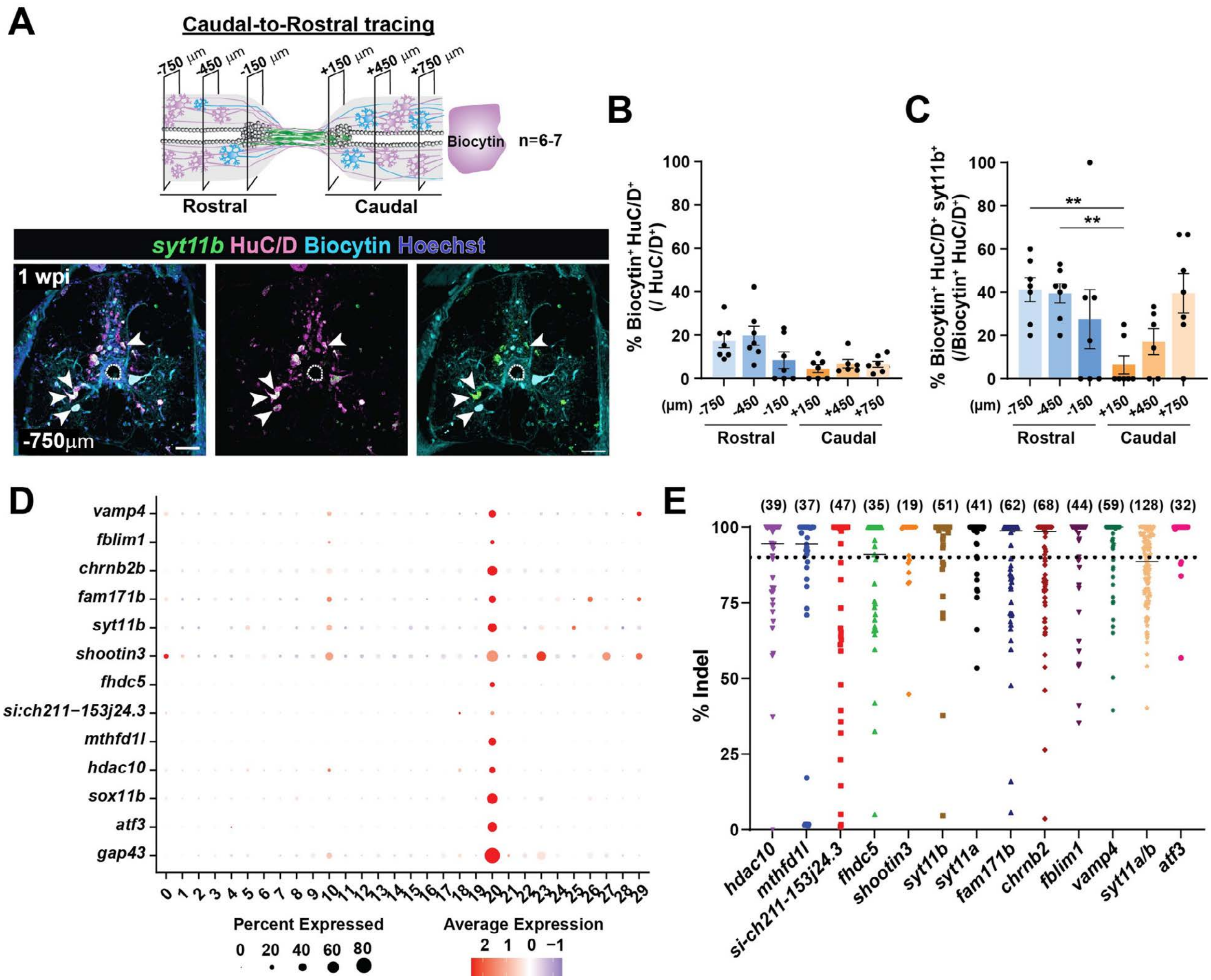

A
Caudal-to-Rostral tracing
-750 µm
-450 µm
-150 µm
+150 µm
+450 µm
+750 µm
Biocytin
n=6-7
Rostral
Caudal
syt11b HuC/D Biocytin Hoechst
1 wpi
-750µm
B
% Biocytin⁺ HuC/D⁺ (/ HuC/D⁺)
(µm) -750 -450 -150 +150 +450 +750
Rostral
Caudal
C
% Biocytin⁺ HuC/D⁺ syt11b⁺ (/Biocytin⁺ HuC/D⁺)
**
**
D
vamp4
fblim1
chrnb2b
fam171b
syt11b
shootin3
fhdc5
si:ch211-153j24.3
mthfd1l
hdac10
sox11b
atf3
gap43
Percent Expressed
0 20 40 60 80
Average Expression
2 1 0 −1
E
(39) (37) (47) (35) (19) (51) (41) (62) (68) (44) (59) (128) (32)
% Indel
hdac10
mthfd1l
si-ch211-153j24.3
fhdc5
shootin3
syt11b
syt11a
fam171b
chrnb2
fblim1
vamp4
syt11a/b
atf3

**Figure S7. CRISPR/Cas9 mutagenesis and regeneration assay of N20 iNeurons (Related to figure 7). (A-C)** Caudal-to-rostral tracing of neurons at 1 wpi. Biocytin was applied 4 mm caudal to the lesion and co-labeled with *syt11b* HCR *in situ* hybridization and HuC/D staining. SC cross sections 750 μm rostral to the lesion are shown in A. Dotted lines delineate central canal edges. The profiles of Biocytin-labeled neurons ($Biocytin^+$ $HuC/D^+$) were normalized to total neurons ($HuC/D^+$) in B. The profiles of Biocytin-labeled neurons ($Biocytin^+$ $HuC/D^+$) were normalized to total neurons ($HuC/D^+$) in B. The profiles of Biocytin-labeled iNeurons ($Biocytin^+$ $HuC/D^+$ $syt11b^+$) were normalized to traced neurons ($Biocytin^+$ $HuC/D^+$) in C. **(D)** Dot plot showing expression of markers selected for mutagenesis in iNeurons. **(E)** Quantification of targeting efficiency using capillary electrophoresis. Percent indel represents the efficiency of indel amplicons relative to wild-type amplicons. For genes in which percent indel efficiency was >90% indel (e.g., *shootin3*), zebrafish were randomly selected fish required for swim endurance assay. For genes with lower targeting efficiency, only genotyped animals with percent >90% were used to assess SC regeneration. Brown-Forsythe and Welch ANOVA test was performed on all four graphs with Dunnett's T3 multiple comparisons test (95% CI). ANOVA p-values are 0.0050 for B and 0.0273 for C. ** $p \leq 0.01$. Scale bars, 50 μm.
